# One-pot lactic acid production from rice straw: A consolidated bioprocess with enzymatic pretreatment-saccharification and Microbial co-fermentation

**DOI:** 10.64898/2026.06.03.729808

**Authors:** Bhavya Surendran V S, Avanthi Althuri

**Affiliations:** Department of Biotechnology, Indian Institute of Technology Hyderabad, Kandi, Sangareddy-502284, Telangana, India

**Keywords:** Green technology, Waste valorization, Co-fermentation, Biorefinery, Sustainability

## Abstract

Global demand for platform chemicals and biomaterials urges us to seek sustainable strategies along with waste valorization to produce lactic acid (LA) sustainably. The study has designed a one-pot fermentation strategy by employing in-house produced ligninolytic and saccharifying enzymes on rice straw along with a consortium of hexose and pentose sugar co-fermenting microorganisms. Biological pretreatment with in-house ligninolytic enzyme was selected for the one-pot strategy from a comparison study of chemical and enzymatic pretreatment of rice straw. In this study, simultaneous pretreatment and saccharification of rice straw followed by LA fermentation by *Lactobacillus casei- Lactobacillus rhamnosus* system (35.58±0.29 g/L) was found out to be more efficient than *Lactobacillus casei-Lactobacillus pentosus system* (29.80±0.92 g/L). Thus, the *L. casei- L. rhamnosus* system (CR system) was selected and was further statistically optimized by response surface methodology (RSM) to yield 64.96 g/L of LA. The fermentation broth was decolorized and purified by ion exchange chromatography to yield 85.56% pure LA with 84.95% optical purity. The one-pot fermentation strategy has reduced the number of unit operations involved to synthesize LA from rice straw without compromising the yield and purity through a greener route. The use of in-house enzymes and consortium of lactic acid producing bacteria in one-pot presents a strategic approach to sustainable LA production. The biological enroute and the minimum use of chemicals during upstream, fermentation, and downstream processing adds to the carbon credit of the process.

**Highlights:**

- Lactic acid was produced from rice straw using one-pot co-fermentation strategy
- Upstream processing employed in-house enzymes from fungal solid-state fermentation
- The process addresses the underutilization of pentose sugars after saccharification
- A consortium LAB produced 64.96 g/L LA with 0.855 g/L.h productivity
- Downstream processing yielded LA with 85.56% purity and 84.95% optical purity

## 1. Introduction

Lactic acid (LA), the three-carbon platform chemical, has a wide variety of uses in pharmaceutical, food, chemical, leather, textile, and cosmetics industries. The major market share of lactic acid is held by cosmetic and pharmaceutical industries. However, there was a surge in LA demand attributed to the use of LA as the raw material for polylactic acid (PLA) synthesis which is a bioplastic having the capacity to replace synthetic plastics and its use in food industry as preservative. Chemical synthesis of lactic acid involves glycerol or acetaldehyde whose origin is petrochemical industry (Tan et al., 2018). These synthesis methods involve extreme reaction conditions, petrochemicals, high energy requirements, and harmful byproducts which influence the overall environmental impact of the process. Climate change and natural resource exhaustion made the governing authorities and stakeholders worldwide encourage the use of bio-based products and reduce the usage of products of non-renewable origin. Biotechnological approaches involve the use of bio-origin renewable materials as raw materials for the synthesis of LA using chemical or biological catalysts under suitable conditions (Wang et al., 2013). In addition to the sustainable aspects of LA fermentation, the specificity of microbial factories to produce optically pure LA has gained attention owing to their involvement in PLA production. The optical purity of LA influences the intrinsic characteristics and biodegradable properties of the polymer synthesized. Certain studies show that chemical catalysts produce both -LA and -LA (Xu et al., 2021; Yang et al., 2026). This racemic mixture can interfere with the efficiency and cost effectiveness of the purification stage. Alternatively, employing selective microbial factories can avoid additional unit operations for optical purification of LA. Lignocellulosic biomass (LCB), categorized as the 2^nd^ generation of feedstock (FS), is non-edible or waste biomass such as crop residues, agri-waste, pruning waste, and forest waste. Its renewable nature makes the process economically viable and environmentally sound fulfilling the sustainability goals. A large fraction of LCB consists of lignin, cellulose, and hemicellulose. These polymeric fractions act as a carbon sink during environmental restoration and as a polymer reservoir during waste valorization. Lignin is responsible for the recalcitrance of the plant structure whereas the other structural polymers; namely, cellulose is made up of hexose sugars while hemicellulose is composed of both hexose and pentose sugars. Since lignin is covalently attached to the carbohydrates in the LCB structure, the presence of lignin in the LCB is a hurdle to achieve economic and energy efficient processes. The intermolecular lignin-carbohydrate bonds respond differently to different cleavage approaches such as alkali, acid, or oxidative methods; hence, the mechanisms and products of reactions may vary. Pretreatment can aid in reducing the recalcitrance of LCB by breaking the lignin-carbohydrate linkages and making structural carbohydrates (cellulose and hemicellulose) available for saccharification to be further broken down into simple sugars. There are several physical, chemical, and biological methods employed to delignify the LCB structure. The chemo-catalytic removal of lignin in LCB uses chemical agents to produce fermentable sugars prior to fermentation with lactic acid producing bacteria (LAB). Yet the presence of microbial inhibitors such as acetic acid and furfural compounds formed during chemical pretreatment can interfere with the saccharification and fermentation processes. Whereas studies that explored biological methods for pretreatment presented a better environment and process-friendly approach. Certain microorganisms like *Paenibacillus alvei*, *Bacillus haynesii* (Rabi Prasad et al., 2024), *Trametes versicolor*, *Phanerochaete chrysosporium*, and *Pleurotus ostreatus* (Tsegaye et al., 2019) are known to produce lignin degrading enzymes in their natural habitat. Generally, lignin degrading enzymes produced by fungi are extracellular and easy to produce (Singh et al., 2023).

The present study explores the potential of agricultural waste, specifically rice straw, to produce lactic acid through a one-pot strategy. This idea of valorizing agricultural waste can solve the dilemma between food security and environmental and economic sustainability. In this consolidated process, the first stage is to prepare the rice straw to expose cellulose and hemicellulose for saccharification. Dilute alkali (sodium hydroxide) and biological (enzymatic pretreatment) treatment were selected as pretreatment methods for rice straw processing. These two approaches were compared through enzymatic saccharification to select an approach with a higher yield of fermentable sugars. A major share of the economics of a process lies in acquiring and processing of chemicals and raw materials involved. The present study aims to make the bioprocess economic by producing the required enzymes; ligninolytic enzymes and saccharifying enzymes in-house and use the extracted cocktail in the process without purification. The fungi *Pleurotus ostreatus* NCIM 1200 and *Trichoderma reesei* NCIM 1186 were used for the production of ligninolytic and saccharifying enzymes, respectively. The fungi *T. reesei* are known to produce both cellulose degrading and hemicellulose degrading extracellular enzymes. This will release both cellulose derived and hemicellulose derived sugars into the saccharifying media. On the contrary to conventional fermentation, one-pot co-fermentation explores the possibility of minimizing the interfaces between upstream processing and fermentation without compromising the efficiency of the process. One-pot co-fermentation of biomass or lignocellulosic biomass refers to a consolidated bioprocessing of pretreatment, biomass hydrolysis, and microbial co-fermentation in a single reactor or vessel. This makes the process less complex and energy-efficient, thereby making it excel in sustainability as well as improving economic competitiveness. This biomanufacturing strategy has shown to reduce the contamination risk and capital and operational costs, eventually improving the process efficiency. The present study envisions to maximize the utilization of monomer sugars during co-fermentation. Saccharification releases a variety of sugars; both hexoses and pentoses. However, there are a few LAB strains recognized to possess both hexose and pentose sugar metabolizing pathways. Hence, attempts have to be made to ensure optimal metabolism to utilize fermentable sugar to form lactic acid in a fermentation system. Co-fermentation approach where a consortium of microorganisms is employed for the production process, makes the process resource efficient and economic. A study by Nancib et al., 2009 reported that a mixed culture of *L. casei* and *L. lactis* utilized 96% glucose and 100% fructose. Whereas the monoculture of *L. casei* utilized 82.2% glucose and 94.4% fructose and *L. lactis* utilized 93.8% glucose and 94.4% fructose (Nancib et al., 2009). The study deduces the possibility of synergistic microbial interactions leading to better substrate utilization and product yield. Hence, the idea of co-fermentation acknowledges the wide spectrum of substrates and metabolic channels that can be employed for product credentials along with reducing the residual sugar and fermentation by-products. This will eventually reduce the carbon footprint of the process by reducing the energy input for waste treatment and enabling complete utilization of available carbon. Therefore, a one-pot co-fermentation approach has been studied to investigate the effect of LAB strains, proportion of the strains in the media, inoculum volume, and duration of fermentation on LA yield. Two different combinations of LAB strains were chosen for the fermentation study; *L. casei: L. pentosus* (CP system) and *L. casei: L. rhamnosus* (CR system) to explore and address the underutilization of hexose and pentose sugars in an LCB saccharified media and produce LA with maximum chiral purity.

## 2. Materials and methods

### 2.1 Raw materials and chemicals

Agri-wastes, rice straw, wheat bran, and coconut coir were collected locally (latitude-17.585286885207577 and longitude-78.11976144726191). Rice straw was oven dried to ensure low moisture content, powdered and sieved using 0.2mm mesh and was characterized for extractives, lignin, and ash using NREL protocol (Sluiter et al., 2008) and Semimicro method was used for cellulose content determination (Updegraff, 1969). Wheat bran washed with water to remove starch and other soluble content was oven dried at 80 until constant weight. Coconut coir was separated from coconut by husking and decorticating and cut into 3 cm length before oven drying at 80 until constant weight. All the chemicals (analytical grade) were purchased from Hyma Synthesis, India.

### 2.2 Microorganisms and growth conditions

The microorganisms used in the present study were procured from the National Collection of Industrial Microorganisms (NCIM), Pune, India and their respective culture media are tabulated in Table 1. LAB strains were grown in MRS broth until ∼10^8^ CFU/ml was achieved prior to inoculating the fermentation media.

**Table 1.** Microorganisms used in one-pot co-fermentation process and their respective culture media.

| S. No | Microorganism | Culture media |
| --- | --- | --- |
| 1 | *Lactobacillus casei* NCIM 5303 | DeMan, Rogosa and Sharpe (MRS) broth media |
| 2 | *Lactobacillus pentosus* NCIM 2912 | DeMan, Rogosa and Sharpe (MRS) broth media |
| 3 | *Lactobacillus rhamnosus* NCIM 5300 | DeMan, Rogosa and Sharpe (MRS) broth media |
| 4 | *Pleurotus ostreatus* NCIM 1200 | Potato Dextrose Agar media |
| 5 | *Trichoderma reesei* NCIM 1186 | Potato Dextrose Agar media |

### 2.3 Production of enzymes

Cellulolytic and xylanolytic enzymes were produced from the solid-state fermentation (SSF) of wheat bran by *T. reesei* (Chakraborty et al., 2016) and the enzyme activities were assayed according to Ghose 2009 (Ghose, 1987). Ligninolytic enzyme was produced by SSF of coconut coir - wheat bran mixture (1:1 ratio) using *P. ostreatus* (Bhattacharya et al., 2011) and the laccase activity was assayed using the guaiacol method (Umar and Ahmed, 2022). After SSF, the extracellular enzymes (both ligninolytic and saccharifying) were extracted from the solid bed using acetate buffer at pH 5 followed by centrifugation at 5000rpm for 10min to remove debris. The clear enzyme cocktails were used without purification for pretreatment and saccharification with appropriate dilution, if needed.

### 2.4 Pretreatment of rice straw

#### 2.4.1 Dilute alkali pretreatment

A modified alkali pretreatment was employed to delignify the rice straw prior to saccharification. Briefly, dilute alkali pretreatment with sodium hydroxide (NaOH) was performed at different alkali loading (20, 60, and 100 mg NaOH/g FS). The weighed feedstock was soaked in alkali solution overnight at room temperature followed by steam explosion at 121°C for 30 min in an autoclave. The solids recovered after the treatment were filtered using muslin cloth, washed till clear filtrate and neutral pH and oven dried at 70 until constant weight for further analysis.

#### 2.4.2 Enzymatic pretreatment

Enzymatic delignification was performed for different enzyme loadings (20, 60, and 100 10^3^ IU/g FS) at 50°C for 24h (Gujjala et al., 2016; Kumar et al., 2017) in a water bath using the ligninolytic enzymes produced from *P. ostreatus*. After the incubation period, the solids were recovered, washed till clear filtrate, and oven dried at 70 for further analysis.

The extent of solids recovery (solid recovery %) was estimated by the gravimetric method. Lignin content in the recovered solids was estimated using NREL method (Sluiter et al., 2008) and the delignification % was estimated using Eq 1.

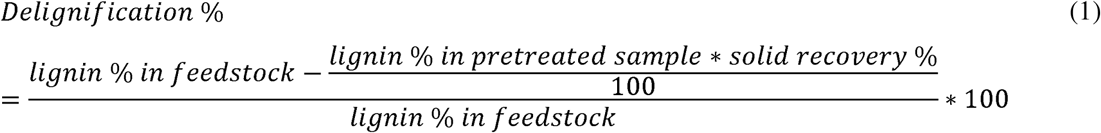

#### 2.4.3 Saccharification of the pretreated sample

The solids recovered from both the pretreatment studies (alkali and enzymatic) were subjected to saccharification using the enzyme cocktail from *T. reesei* culture for 24h at 50°C in a water bath (Avanthi and Banerjee, 2016, 2016) with 10%(w/v) solid loading to compare the pretreatment efficiency. The reducing sugars produced during saccharification were estimated by the dinitro salicylic acid (DNS) method (Miller, 1959). The pretreated sample, which showed better hydrolysis, was selected for further studies and the compositional and structural changes in the pretreated rice straw were analyzed by Fourier Transform Infrared (FTIR) Spectrometry and X-Ray Diffractometry. FT-IR spectroscopy of untreated rice straw (control) and enzyme treated rice straw samples were run from 4000-400 cm^-1^ (Make: Jasco, Model: FTIR-4600). The XRD patterns were obtained using λ = 0.154 nm radiation source (PANalytical Pro) and the crystallinity index (CrI) was estimated using Eq 2 (Phitsuwan et al., 2017).

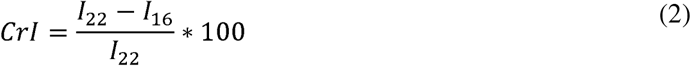

Where, I_22_ is the intensity of peak at 2θ=22 and I_16_ is the intensity of peak at 2θ=16 obtained from the XRD pattern.

### 2.5 Comparison between sequential pretreatment and saccharification (SqPS) and simultaneous pretreatment and saccharification (SPS)

The two stages involved, namely pretreatment and saccharification, are optimized to improve product yield along with productivity. The conventional strategy is to pretreat the LCB followed by saccharification. A one-pot approach at the saccharification stage calls for stability of the chemical and/or biochemical catalysts and prevention of product or by-product inhibition in a set-up where pretreatment and saccharification happen sequentially or simultaneously. With the aim of making the whole bioprocess consolidated, shorter, and simpler, a comparison study was drawn between sequential enzymatic pretreatment and saccharification (SqPS) and simultaneous enzymatic pretreatment and saccharification (SPS). In contrast to the conventional flow of unit operations, ligninolytic enzymes and saccharifying enzymes were mixed in 1:1 ratio with 10%(w/v) feedstock for SPS making the final enzyme loadings to be 5000 IU/g laccase, 45 IU/g carboxymethyl cellulase (CMCase), 75 IU/g xylanase, and 10.5 FPU/g FPase. The SPS reaction mixture was incubated 24h at 50°C in a water bath. In SqPS, 10%(w/v) feedstock was incubated in a water bath at 50°C for 8h in delignifying enzyme followed by the addition of equal amount of saccharifying enzyme and incubated at 50°C for 24h. The supernatant after saccharification was obtained by centrifugation at 5000 rpm for 30 min and the RS sugar content was estimated by the DNS method.

### 2.6 One pot co-fermentation to produce lactic acid

Fermentation systems were studied for maximum LA production using hexose-utilizing and pentose-utilizing LAB. The sugars produced from pretreatment and saccharification were subjected to LA fermentation by two different systems: *L. casei* and *L. pentosus* (CP) system and *L. casei* and *L. rhamnosus* (CR) system. Both these systems were evaluated by one variable at a time (OVAT) approach, and the one resulting in higher RS concentration and LA (after purification) yield was selected for further studies. All the fermentation experiments were performed at 10ml culture media capacity in batch fermentations mode.

#### 2.6.1 Ligninolytic: saccharifying enzyme ratio

The effect of ligninolytic (laccase): saccharifying (cellulases plus xylanase) enzyme ratio in the production of fermentable sugars from rice straw was studied by varying the amount of ligninolytic enzyme in the total enzyme cocktail in terms of fraction of ligninolytic enzyme. At 10%(w/v) rice straw loading, a varying volume of ligninolytic enzyme (0, 1, 2, 3, 4, 5, and 6ml) and rest of saccharifying enzyme, for a total of 10ml working volume was added and incubated for 24h at 50°C in the water bath. The liquid was separated after incubation by centrifugation at 5000rpm for 30 min and the reducing sugar content of the clear supernatant was estimated using the DNS method. The ligninolytic: saccharifying enzyme ratio resulting in higher RS yield was selected for subsequent studies.

#### 2.6.2 Feedstock concentration

A varying concentration of rice straw (10, 15, 20, and 25%w/v) was added to 10 ml of ligninolytic-saccharifying enzyme cocktail and incubated for 24h at 50°C in a water bath. The supernatant was collected and the reducing sugar (RS) concentration was estimated by the DNS method (Miller, 1959). Rice straw concentration resulting in higher RS yield was selected for further studies.

#### 2.6.3 SPS reaction time

Pretreatment and saccharification was conducted with the optimized ligninolytic enzyme concentration and feedstock concentration at 50°C in water bath for different SPS reaction times with 6h intervals (0, 6, 12, 18, 24, 30, 42, and 48h). After incubation the supernatant was collected by centrifugation at 5000 rpm for 30 min and the reducing sugar concentration was estimated by DNS method.

#### 2.6.4 LA co-fermenting systems

The above selected OVAT conditions were used to conduct the SPS of rice straw. The treated rice straw slurry was then supplemented with the following composition of supplements prior to fermentation: MgSO_4_.7H_2_O (0.8g/L), K_2_HPO_4_ (1.2g/L), KH_2_PO_4_ (1.2g/L), Yeast extract (2g/L), and Tween 20 (0.4g/L). The CP and CR systems were studied separately during the co-fermentation process. The fermentation experiments were conducted at 30°C and 120 rpm. Parameters such as co-fermentation time (0-108h), internal ratio (1:0, 1:3, 1:1, 3:1, and 0:1) of LAB strains, and inoculum volume (6, 8, 10, 12, 14 v/v%) were optimized using the one variable at a time (OVAT) approach.

#### 2.6.5 Decolorization of fermentation broth

Powdered activated carbon (PAC) was used to decolorize the fermentation broth. After fermentation at the optimized conditions the broth was centrifuged at 5000rpm for 15min to separate the broth and residual solids. The pH of the broth was adjusted to 2.0 using dil. HCl to precipitate the soluble proteins in the broth. Different concentrations of PAC (5, 10, and15 w/v%) were studied to probe for the best concentration to decolorize the broth (de Oliveira et al., 2021). Definite amount of PAC was added to the broth and incubated at room temperature for 30min at 100rpm followed by centrifugation to remove the solid residue. The decolored liquid obtained was filtered through a 0.45μm cellulose acetate filter and used for further purification.

#### 2.6.6 LA purification by chromatography

A weak anion exchange resin, Amberlite^®^IRA-67 was selected for LA purification (Ahmad et al., 2021). The resin was washed with 1M HCl and 1M NaOH followed by water wash to convert the free base form into Cl^-^ form. These converted resins were oven-dried at 50 and stored in airtight container to prevent further moisture adsorption before chromatography use. The purification of decolored fermentation broth for both CP and CR systems was conducted at 25 with 1N HCl as eluent (1ml/min flow rate) in a column with 2.5 mm diameter and 45 cm length. The LA concentration in decolored broth was maintained at 25g/L and pH 3.0. After elution, the resin was regenerated using 1N NaOH followed by water wash. The concentration and purity of LA in the eluted samples was determined using HPLC.

### 2.7 Optimization of LA production by sequential co-fermentation using RSM

The best performing co-fermentation system amongst CP and CR from OVAT studies was selected in this study for further optimization of LA production. One-pot co-fermentation system was mathematically modeled using Response Surface Methodology (RSM) using the software MINITAB^®^ 22. RSM models the system in terms of a second-order polynomial which showcases the individual effects of input variables along with the effect of interaction of one over the other on LA production. The co-fermentation stage after simultaneous pretreatment and saccharification was statistically optimized by central composite design (CCD) using four independent variables at five levels with 16 cube points and 8 axial points. Fraction of *L. casei* in the consortium (%), inoculum volume (v/v%), time interval between *L. casei* and *L. rhamnosus* inoculation (h), and co-fermentation time after *L. rhamnosus* inoculation (h) were selected as the independent variables. Each parameter was studied at 5 levels (-α, −1, 0, +1 & + α) with α = 2 in the parameter ranges obtained from the OVAT study and are tabulated in Table 2.

**Table 2.**
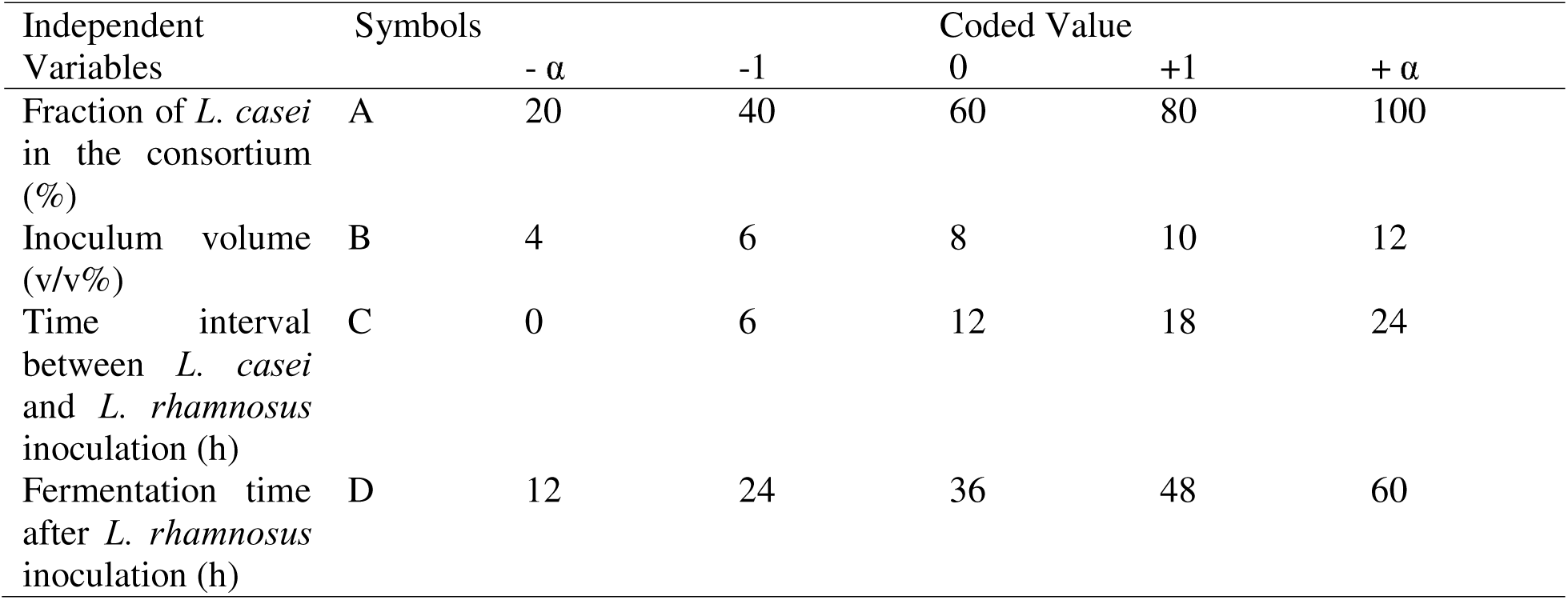
Independent variables and their coded values used in CCD based RSM.

| Independent Variables | Symbols | Coded Value |  |  |  |  |
| --- | --- | --- | --- | --- | --- | --- |
| | | $-\alpha$ | -1 | 0 | +1 | $+\alpha$ |
| Fraction of *L. casei* in the consortium (%) | A | 20 | 40 | 60 | 80 | 100 |
| Inoculum volume (v/v%) | B | 4 | 6 | 8 | 10 | 12 |
| Time interval between *L. casei* and *L. rhamnosus* inoculation (h) | C | 0 | 6 | 12 | 18 | 24 |
| Fermentation time after *L. rhamnosus* inoculation (h) | D | 12 | 24 | 36 | 48 | 60 |

The design of experiments (DOE) obtained (30 experimental runs) were performed experimentally in duplicates, where the conditions of simultaneous pretreatment and saccharification were adopted from the previously conducted OVAT studies. Experiments were conducted for a reaction volume of 10ml and 5% CaCO_3_ was added to the SPS liquid media before sterilizing the contents at 121□ and 15 psi for 15 min. After co-fermentation the LA content was estimated by FeCl_3_ method (Borshchevskaya et al., 2016). Based on the data obtained from the experimental runs, the system was modelled by regression analysis in terms of the independent variables (X) and the response, LA yield (Y) using a second-order polynomial equation (Eq 3).

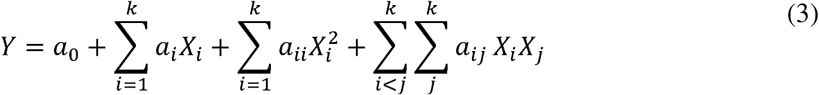

Where Y is the predicted response (LA concentration), k is the number of independent variables evaluated, X_i_ are the independent variables, and a_0_ are the regression coefficients. The quadratic equation generated was optimized for maximum LA yield, and a set of validation experiments were conducted to verify the model. The ANOVA and F-test were conducted to evaluate the significance of each independent variable on the response. The coefficient of determination R^2^ was determined to express the quality of fit of the quadratic equation onto the experimental data. Three-dimensional interaction plots were obtained for two independent variable sets to study the interaction between them and their respective effect on the response, LA concentration.

LA was produced at optimum conditions predicted by the RSM method and purified as depicted in Fig 1. Briefly, the broth after co-fermentation was separated and acidulated with H_2_SO_4_ to convert calcium lactate to lactic acid. This was followed by decolorization at the optimum condition obtained for section 2.5.5 and was purified by chromatography as detailed in section 2.6. The samples were characterized for their purity by HPLC analysis, and the optical purity of the purified sample was determined by CD Spectroscopy.

**Fig 1.**
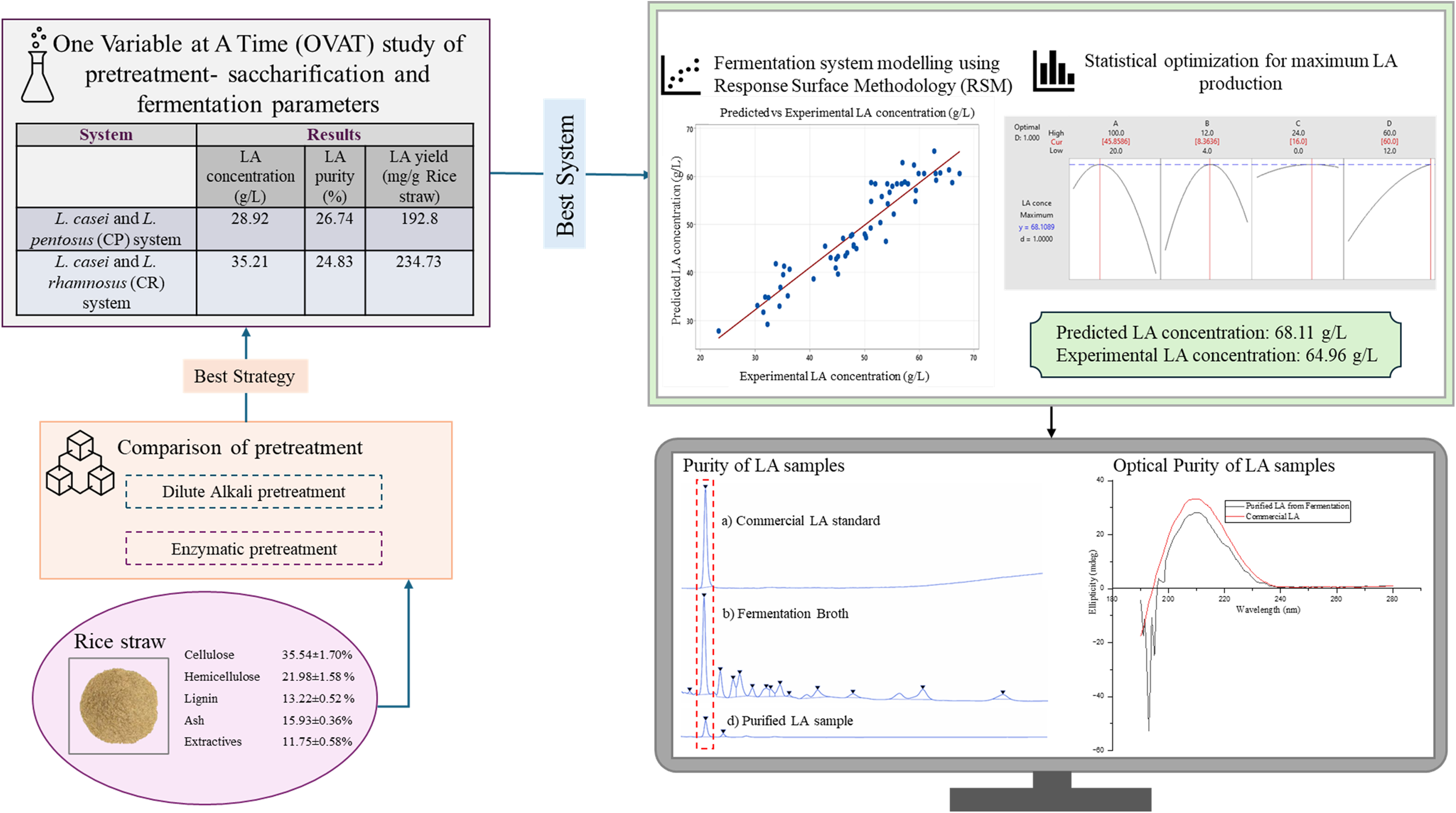
Holistic view of the LA production and purification process from rice straw by one-pot co-fermentation strategy

### 2.8 Characterization

The stereospecificity of LA in the purified sample was calculated using Eq 4 in comparison to commercial L-LA (100% purity) and thereby the optical purity of the sample was determined. The optical rotation of the sample was analyzed using circular dichroism spectropolarimeter

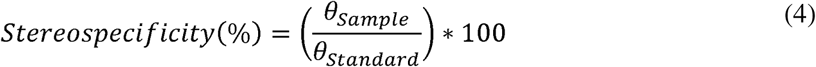

### 2.9 Analytical methods

The concentration of reducing sugars was estimated using the DNS method (Miller, 1959) and lactic acid concentration by iron (□) chloride method (Borshchevskaya et al., 2016). The purity and LA concentration of chromatographic elutions was estimated using high performance liquid chromatography (HPLC) (Agilent 1200 infinity series, Santa Clara, CA, USA) equipped with C18 column (300 mm × 7.8 mm; Bio-Rad Laboratories Inc., Hercules, CA, USA) and Diode Array Detector (DAD) at 210 nm. The mobile phase was 20mM phosphate buffer (pH 3.0) with 1.5ml/min flow rate for a run time of 10 min and injection volume of 5 μl (Kishore et al., 2013). Hexose and pentose sugars in the fermented broth were analyzed by the HPLC method using Shim-pack GIST C18 (5 μm) separation column (250 × 4.6 mm) and DAD at 248 nm. The Shimadzu HPLC system was used with 7.7g/L ammonium acetate solution as mobile phase A and 100% acetonitrile as mobile phase B. The sugar samples for analysis were prepared by modifying the method reported by Wang et al (Wang et al., 2021). Briefly, 0.5ml of fermented broth was mixed with an equal volume of 0.3M NaOH solution followed by the addition of 1ml of 0.5M 1-phenyl-3-methyl-5-pyrazolone (PMP) - methanol solution and vigorously mixed for 1min. Further, the mixture was incubated at 70 for 120min. After incubation, the reaction mixture was neutralized by adding 1ml of 0.3M HCl followed by the addition of 5ml chloroform to remove the unreacted PMP. The upper layer of the reaction mixture was extracted and filtered using a 0.22μm filter for HPLC analysis. A gradient system mobile phases (0 min; B-20%, 20 min; B-30%, and 30 min; B-20%) was employed for a run time of 30 min with flow rate 0.8ml/min and 10µL injection volume. The samples and mobile phases were filtered through 0.22μm filters before HPLC analysis. Statistical significance of each experiment was estimated using analysis of variance (ANOVA) and the confidence was set at p<0.05.

## 3. Results and discussion

### 3.1 Feedstock Composition and In-house Lignocellulolytic Enzymes

The composition of rice straw was evaluated using the NREL method to assess its potential for lactic acid production. The study showed that rice straw contains 35.54±1.70% cellulose, 21.98±1.58 % hemicellulose, and 13.22±0.52 % lignin along with 15.93±0.36% ash and 11.75±0.58% extractives on dry mass basis. The cellulose, hemicellulose, and lignin contents are in similar ranges of reported case studies. Jin and Chen., 2017 reported a composition of 33.9±2.8% cellulose, 25.6±3.3% hemicellulose, and 10.2±1.6% Klason lignin in the rice straw (Jin and Chen, 2007). Whereas Raudhatussyarifah et al., 2022 reported the rice straw to contain 37.5% cellulose, 35.5 % hemicellulose, and 14.5 % lignin (Raudhatussyarifah et al., 2022). The difference in the composition can be attributed to geographical, cultivational, and seasonal variations. The activity of ligninolytic enzymes produced from *P. ostreatus* SSF on coir-wheat bran was found to be 1000 IU/ml. Whereas, enzyme activity of CMCase, xylanase and FPase was 9 IU/ml, 15 IU/ml, and 2.1 FPU/ml, respectively in *T. reesei* wheat bran SSF cocktail.

### 3.2 Alkali vs. Enzymatic Pretreatment of Rice Straw

The delignification %(w/w) and solid recovery %(w/w) from alkali and enzymatic pretreatment are represented in Fig 2a. In case of chemical pretreatment with NaOH, it was observed that as the alkali loading increased, delignification in the feedstock increased. The data showed that 100mg NaOH/g FS corresponding to 1.0% NaOH concentration for 10% feedstock loading was able to remove 75.10±3.09% (w/w) of total lignin from the feedstock. Theoretically, the method should retain 90.06% of total solids; however, the pretreatment retained only 45.94±1.09% (w/w) solids. This indicates that in addition to removing the lignin from the feedstock, alkali pretreatment removes other structural components including carbohydrates such as cellulose and hemicellulose from the feedstock. The increase in delignification% of rice straw is in accordance with the reported study (Ningthoujam et al., 2023). As per their study, 4% NaOH treatment reduced the lignin content from 17% to 4% while the relative cellulose content increased from 29% to 45%. Whereas in the present study, 100mg NaOH/g FS treatment reduced the lignin content from 13.22% to 7.17% while relative cellulose content increased from 35.54% to 47.58%. Despite the promising delignification and solid recovery results, the need to neutralize the biomass after alkali pretreatment by water wash poses a problem at larger scales, since it can severely affect the overall cost of production.

**Fig 2.**
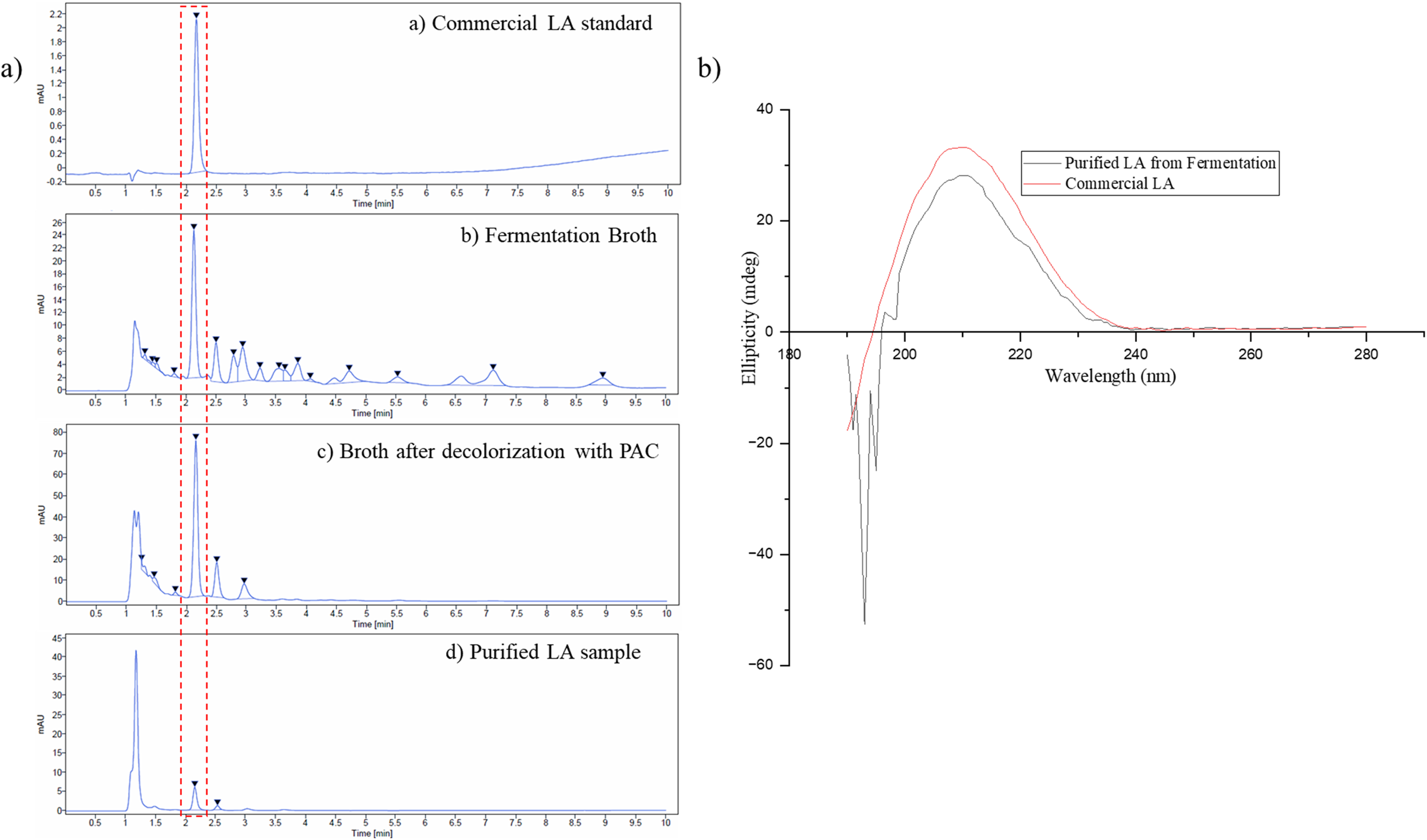
Pretreatment comparison: a) Solid recovery % and delignification % of alkali pretreatment and enzymatic pretreatment and b) RS concentration (g/L) and RS yield (g/g FS) of alkali pretreatment and enzymatic treatment (p<0.05).

The enzymatic pretreatment followed similar trends of solid recovery and delignification as in alkali pretreatment, however, with higher delignification% and solid recovery%. The pretreatment efficiency upon saccharification was investigated by saccharifying all the solid residues recovered after both alkali and enzymatic pretreatment (Fig 2b). It was found that enzymatic pretreatment resulted in higher RS concentration and RS yield as compared to alkali pretreatment. The difference in RS concentration can be attributed to the higher delignification% and better accessibility of cellulose and hemicellulose in the enzyme pretreated rice straw to saccharifying enzymes. Whereas the difference in RS yield is due to the differences in solid recovery % wherein NaOH treatment caused higher loss of structural carbohydrates than enzymatic pretreatment. These structural changes were reported to affect the saccharification efficiency. However, Van Dyk and Pletschke., 2012 reported that carbohydrate binding domains (CBM) of cellulase is responsible for the precise binding of the enzyme to the substrate and the sensitivity of these domains affect the efficiency of saccharification and thereby the rate of reaction (Van Dyk and Pletschke, 2012). It was stated that some CBMs bind to the crystalline regions of cellulose in the substrate and stabilize the enzyme on the substrate surface thereby aiding in the hydrolysis process (BORASTON et al., 2004). In the present study, the rationale for alkali pretreatment resulted in lesser RS yield is the loss of cellulose along with lignin from rice straw which influences the binding of CBMs. Whereas the enzymatic pretreatment gave a suitable environment for CBMs binding. Hence, the saccharification efficiency can be drawn as a function of the type of pretreatment employed. In addition, Pallapolu et al., 2011 reported that in switch grass lime pretreatment with 51.87% delignification showed higher cellulose digestibility than ammonia pretreatment with 61.21% (Pallapolu et al., 2011). The study infers that the surface modification during pretreatment is as crucial as delignification% in lignocellulosic biomass (Rollin et al., 2011). Since enzymatic pretreatment resulted in better delignification and saccharification efficiency, it was selected for future studies.

#### 3.2.1 Characterization of ligninolytic enzyme treated rice straw

To gain a better insight into the structural changes during enzymatic pretreatment, before and after treatment samples of rice straw were studied through FT-IR and XRD analyses. The FT-IR spectrum of the untreated and pretreated rice straw (Fig 3a) shows the evidence of lignin removal from the structure. Reduction in intensity of hydroxyl groups of aliphatic and phenolic groups of lignin (3500-3400 cm^-1^), -CH stretching of methyl, methylene and methoxyl side groups of lignin (2910-2840 cm^-1^) (Boeriu et al., 2004), aromatic ring structure in lignin (1648, 1512 and 1427 cm ¹), and conjugated carbonyl (C=O) group in lignin (1659 cm ¹) showed the removal of lignin by enzymatic pretreatment compared to the untreated rice straw. The presence of cellulose was confirmed by the CH_2_ vibration at 898 cm^-1^ and C-O-C vibration at 1430 cm^-1^ corresponding to crystalline and amorphous regions of cellulose, respectively (Jin et al., 2020) While the shoulder peak at 1736 cm ¹ is associated with the carbonyl band (C=O) in hemicellulose proving the presence of hemicellulose in the pretreated rice straw (Zhang et al., 2018).

**Fig 3.**
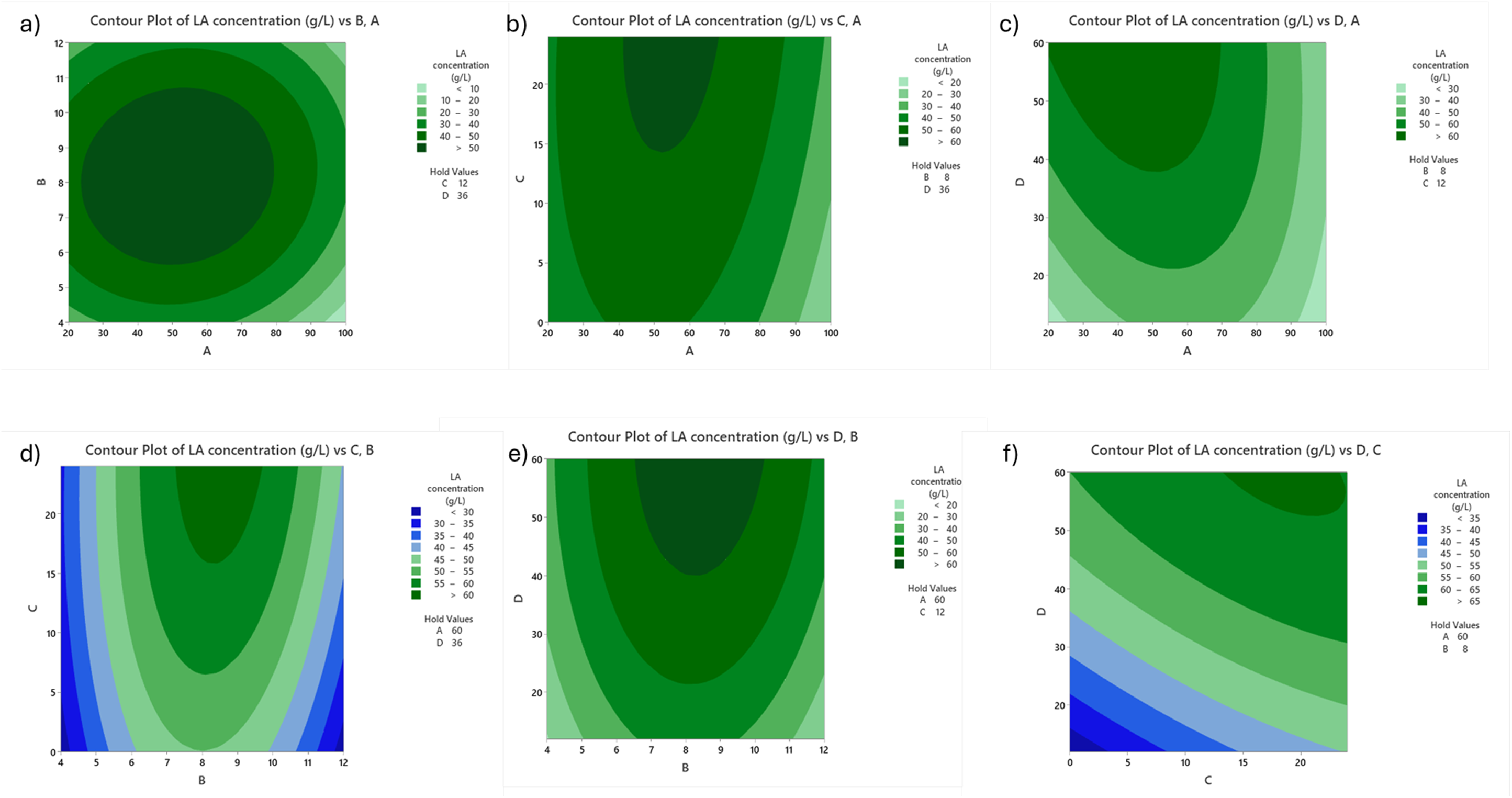
Comparison of rice straw (black line) and enzyme treated rice straw (red line) by structural characterization: a) FTIR spectra and b) XRD spectra

XRD analysis reveals structural changes owing to enzymatic pretreatment suggesting delignification. The XRD spectra of untreated and pretreated rice straw is exhibited in Fig 3b. The increase in the intensity of peak at 22° and 16° indicates the pretreatment has revealed the crystalline cellulose in the structure (Moneim et al., 2018; Ningthoujam et al., 2023). Moreover, the decrease of peak at 18.3° in the treated rice straw reveals that the amorphous region of cellulose has lessened upon pretreatment. The crystallinity index (CrI) of rice straw sample was found to be 69.48% and that of enzyme treated rice straw was 74.99% which underscores the removal of amorphous regions from rice straw structure. Hence, it can be concluded that the enzymatic pretreatment has exfoliated the rice straw structure by stripping lignin to reveal the structural carbohydrates. Ningthoujam et al 2023 reported that the increased crystallinity of rice straw structure facilitates enzymatic saccharification (Ningthoujam et al., 2023). Similar observations were reported by Phitsuwan P et al., 2017, where laccase pretreatment has resulted in the removal of lignin and partial degradation of hemicellulose exposing the crystalline cellulose. This increase in crystallinity reduced the nonproductive absorption of enzyme, eventually leading to higher reducing sugar content (Phitsuwan et al., 2017).

### 3.3 Comparison between SqPS and SPS

The initial experiments on the efficiency of alkali and enzymatic pretreatment revealed that enzymatic pretreatment is efficient, thus further studies on sequential (SqPS) and simultaneous (SPS) pretreatment and saccharification were designed using enzymes. The comparison study showed RS productivity of 2.64±0.08 (g/L.h) from SqPS (0.681±0.019 g/g FS) and 3.01±0.04 (g/L.h) from SPS (0.581±0.009 g/g FS). The productivity data evidently supports simultaneous pretreatment and saccharification (SPS) as a relatively rapid process considering that the aim is to design a simpler and shorter one-pot fermentation strategy for LA production. In SPS, delignifying enzymes depolymerize lignin and consequently improve the cell wall permeability of rice straw. This facilitates cellulases and xylanases to access cellulose and xylan, respectively for improved saccharification. Both the category of enzymes used in SPS; delignifying and saccharifying were found to be active in the temperature range of 45-55 and pH range of 4.5-5.5 making it possible to combine both the unit operations. Since ligninolytic and saccharifying enzymes were acting synergistically in the SPS reaction mixture, reaction rate improved and accordingly productivity was enhanced. Balasubramaniam and Rajarathinam 2013 reported that when rice straw was biologically pretreated with *P. ostreatus* for 44 days followed by cellulase mediated saccharification, it produced ∼80 g/L glucose in the first 24h of incubation and later reached to ∼100g/L glucose within 35h of incubation for a cellulase loading of 25 FPU/g FS (Balasubramaniam and Rajarathinam, 2013). A similar trend for reducing sugar concentration was observed in the present study with 10.5 FPU/g FS; ∼80 g/L RS concentration for 18h and ∼80 g/L for 24h which can be attributed to synchrony in enzyme action. Moreover, SPS reduced the number of unit operations along with the reaction time relative to SqPS. This positively impacts overall productivity and improves the process economy. Thus, the strategy of SPS was selected for further studies. Masran et al., 2020 reported trends where SPS resulted in higher productivity (0.151 g/L.h) in comparison to SqPS (0.061 g/L.h) reducing the overall process time and improving the resource recovery from the biomass (Masran et al., 2020).

### 3.4 OVAT study of one-pot co-fermentation process

Optimization of process variables was initiated with OVAT study for maximizing RS and LA production. The optimization procedure is divided into three stages: simultaneous pretreatment and saccharification stage, co-fermentation stage, and decolorization and purification stage. The effect of process variables on SPS and co-fermentation was evaluated, and the results are depicted in Fig 4.

**Fig 4.**
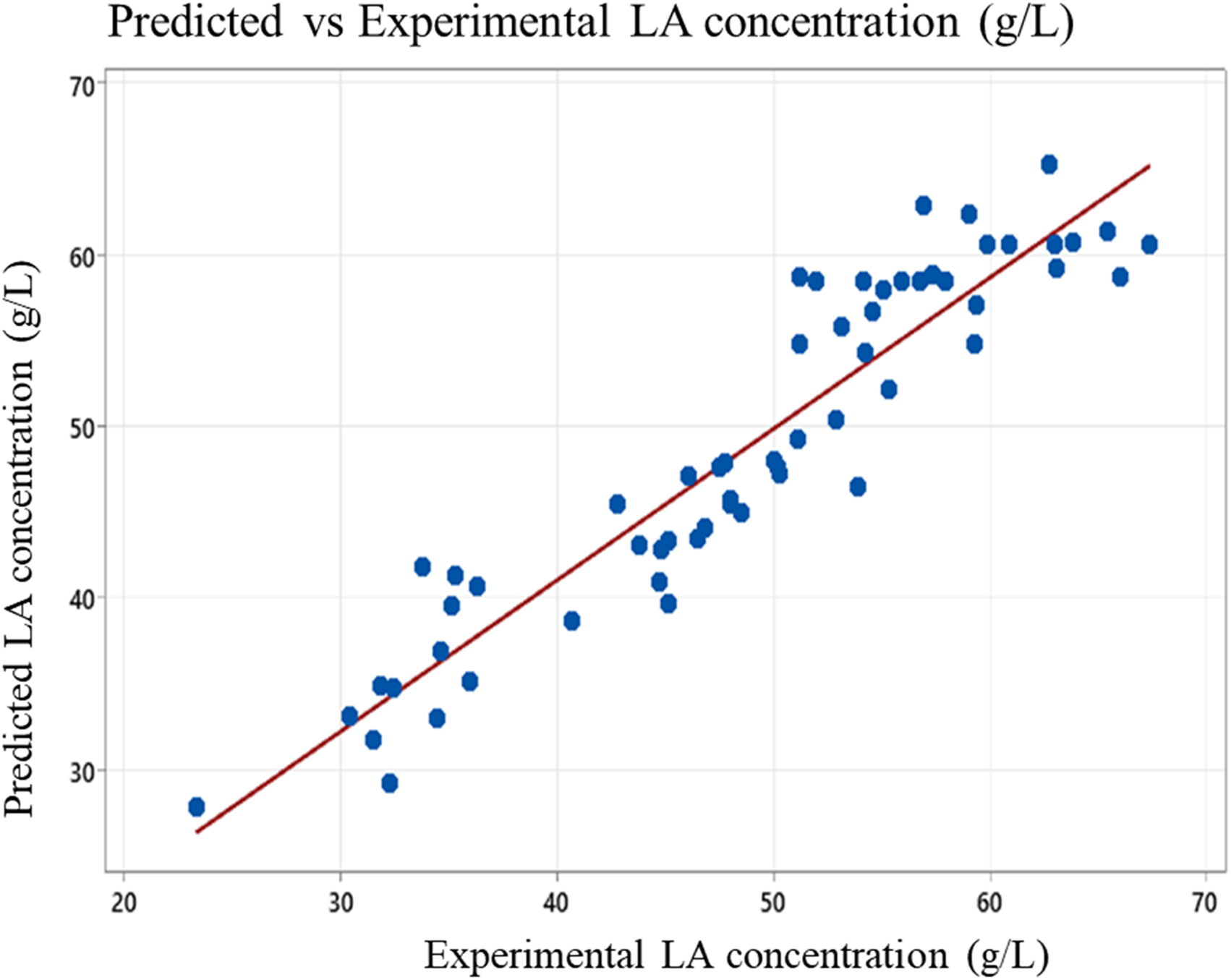
Influence of process variables on the production of reducing sugars through SPS and LA through subsequent co-fermentation using both CP and CR systems (p<0.05): a) Effect of ligninolytic enzyme content, b) Effect of feedstock concentration, c)Effect of SPS reaction time, d) Effect of internal ratio of LAB strains and co-fermentation time (black line-RS concentration and red line-LA concentration), and e) Effect of inoculum volume f) Effect of PAC dosage on the decolorization of fermentation broth

#### 3.4.1 Simultaneous Pretreatment and Saccharification

For optimizing the process variables of the SPS process, ligninolytic enzyme cocktail from *P. ostreatus* and saccharifying enzyme cocktail from *T. reesei* was employed. The process variables were evaluated based on RS production in the broth.

##### 3.4.1.1 Effect of ligninolytic: saccharifying enzyme ratio

During the SPS process, delignifying enzymes start the depolymerization of lignin, and the process of hydrolysis is conducted by cellulases and xylanases on cellulose and xylan, respectively. The ratio of ligninolytic and saccharifying enzyme required for SPS depends on the biochemical composition of rice straw. From Fig 4a, it can be observed that RS concentration continued to increase till 40% ligninolytic enzymes and then decreased. The decrease in RS concentration beyond 40% can be due to the overcrowding and stearic hindrance between saccharifying enzymes and ligninolytic enzymes. The overcrowding of excess saccharifying enzymes mask the active sites for ligninolytic enzymes and thereby interfere with lignin removal, which consequently results in low fermentable sugars (Masran et al., 2020). Therefore, a balance between ligninolytic and saccharifying enzymes is vital in directing cellulases towards cellulose-active sites. Cellulases are known to form non-productive linkages with lignin which can in turn reduce the saccharification efficiency and cellulase recyclability (Moilanen et al., 2011; Rahikainen et al., 2013). Thus, cellulose degradation in the saccharification process is a synergistic effect of laccases and cellulases. This synergy is a function of rates of reaction, type of products formed, properties of substrate, and experimental conditions. The present evaluation showed the highest synergistic effects when 40% of the concoction is made up of ligninolytic enzyme from *P. ostreatus* SSF.

##### 3.4.1.2 Effect of feedstock concentration

The impact of feedstock concentration on the production of fermentable sugars by saccharifying enzyme cocktails is shown in Fig 4b. It can be inferred that the concentration of RS increased drastically until 15% FS and then there was no significant increase in RS concentration. High amounts of feedstock resulted in diffusion-limited, highly viscous system due to high solid and low liquid content. This in turn reduces the enzyme units available for unit grams of FS and increases the apparent viscosity of the mixture, making the handling and mixing difficult. In addition, the product (reducing sugar) concentration tends to increase as the substrate concentration increases, eventually leading up to product inhibition of the enzymes (Modenbach and Nokes, 2013). The cumulative effect of increased product inhibition and reduced mass transfer rate at high feedstock concentration does not favor better saccharification. This can be the rationale for the saturation of reducing sugar concentration at higher FS concentrations. To conclude, an optimum concentration of feedstock represents a system where the substrate, product, and liquid work complementarily. Hence, 15% FS concentration corresponding to cellulase enzyme loading of 8.35 FPU/g rice straw was selected for further studies. Similar pattern of results was reported by Wood et al., 2016 where they found that the optimum cellulase (Cellic^®^ CTec2 enzyme) dosage for saccharification was 6.49 FPU/g rice straw (Wood et al., 2016). Where they reported a sugar yield (in terms of glucose) of ∼90g/L which is comparable to the present study.

##### 3.4.1.3 Effect of SPS reaction time

The kinetics of enzymatic saccharification of enzyme pretreated rice straw from 0-42 h is shown in Fig 4c. It was observed that the enzyme activity and hence RS concentration have risen in the first 18h of duration and slowed down after 24h. However, the trend shows that after 18h of incubation, there is only marginal improvement in the saccharification process. The decrease in enzyme activity can be due to the inhibitors produced during hydrolysis (Xu et al., 2007). Similar results were reported by Marques NP et al., 2018, where maximum saccharification was achieved by the end of 20h of saccharification using enzyme cocktails from *Botryosphaeria sp.* and *Saccharicola sp.* on sugarcane bagasse feedstock (Marques et al., 2018). Whereas, dos Santos NK et al 2022 observed that saccharification efficiency is a function of the synergistic effects of the enzymes involved when enzyme cocktails are used. In the case of simultaneous pretreatment and saccharification, as the lignin was getting hydrolyzed, the cellulose and hemicellulose becomes available for holocellulases, as a result the concentration of reducing sugars increases in the broth improving the productivity of the process (Pasquini, 2025). It can be concluded that the use of in-house saccharifying enzyme cocktails did not compromise the productivity of the system compared to the reported studies conducted with commercial enzymes.

#### 3.4.2 Fermentation of saccharified rice straw

##### 3.4.2.1 Effects of internal ratio of LAB strains and co-fermentation time

The lactobacillus bacterial strains *L. casei*, *L. pentosus*, and *L. rhamnosus* were cultured in the enzyme treated rice straw broth and the concentrations of reducing sugars (RS) and lactic acid (LA) were monitored. The trend of formation of lactic acid was studied along with the reduction of reducing sugars during fermentation. Fig 4d shows the pattern of reducing sugar utilization and lactic acid production from LAB strain concoctions and pure cultures. It was observed that *L. rhamnosus* produced slightly more lactic acid than *L. pentosus* and *L. casei* produced least amount of LA. Since glucose and xylose are the dominant sugars produced from structural carbohydrates of rice straw, the selection of microorganism has to be made to utilize both hexose and pentose sugars to their maximum. There are different pathways of producing LA by a microorganism: homofermentative pathway and heterofermentative pathway. Most LAB are known to have homolactic acid pathway and they exclusively produce LA using hexose sugars. Whereas, there are some LAB which utilize both hexose and pentose sugars via heterolactic acid pathway to produce LA. However, the by-products such as acetic acid and ethanol are the drawbacks of heterolactic pathway. Most of the heterolactic acid producing microorganisms exhibit a hierarchical consumption pattern for sugars such that they tend to consume xylose only in the absence of glucose. It was observed that *L. casei* utilized 27.98% of total reducing sugars initially present in the fermentation media. However, *L. pentosus* and *L. rhamnosus* utilized 32.86% and 39.46% of total reducing sugars. This can be interpreted along with the efficiency of *L. pentosus* and *L. rhamnosus* to ferment both glucose and xylose whereas *L. casei* to ferment only glucose (Jin et al., 2020; Xu et al., 2007). This explains the variation in reducing sugar utilization between the hexose fermenting *L. casei* and hexose and pentose fermenting *L. pentosus* and *L. rhamnosus.* However, the heterolactic pathway of *L. pentosus* and *L. rhamnosus* along with product inhibition by lactic acid seemed to impact the LA production. Hence, the LA concentrations of the pure culture fermentation were comparable; *L. casei*-27.84±0.65 g/L, *L. pentosus*-28.19±0.72 g/L, and *L. rhamnosus*-29.22±0.41 g/L. From Fig 4d, it can be observed that *L. casei* and *L. pentosus* achieved saturation in LA concentration within 48-60h fermentation time, whereas *L. rhamnosus* took 72-84h to achieve saturation, indicating relatively high tolerance of *L. rhamnosus* to LA concentration (Hudeckova et al., 2018). Similarly, the yield of LA on reducing sugar was 0.355 g/g for *L. casei*, 0.360 g/g *L. pentosus*, and 0.378 g/g for *L. rhamnosus*, respectively. Similar results were reported for certain monoculture studies. Costa et al., 2022, for example, reported that *L. casei* produced 22.06 g/L LA with a yield of 0.48 g/g sugar from cheese whey (Costa et al., 2024). Whereas, Chen et al., 2020 reported 31.0 g/L LA produced from liquefied cassava bagasse using *L. rhamnosus* with a yield of 0.94 g LA/g glucose (Chen et al., 2020) and Hu et al., 2016 reported 33.7g/L LA production with a yield of 0.42 g/g from alkali treated corn stover using *L. pentosus* (Hu et al., 2016).

In natural habitat, co-existence of microbial strains is common and is beneficial for certain metabolite consumption and/or production. Thus, co-fermentation of microorganisms has gained interest from researchers in the synthesis of valuable chemicals. The co-fermentation studies with two different combinations of LAB (CP and CR systems) showed better yield of LA from rice straw than fermentations with individual microbial strains. The effect of internal ratio of LAB strains in the co-fermentation of saccharified rice straw is shown in Fig 4e. The effect of inoculation proportion in terms of internal ratio shows the strain interactions in the fermentation broth. Both the pentose sugar utilizing LAB strains had added higher LA in the broth along with improved sugar utilization. It was observed that in case of co-fermentation with *L. casei* and *L. pentosus* equal volume of both the strains produced maximum amount of lactic acid (30.37±0.56g/L) with a productivity of 0.502 g/L.h. The amount of *L. pentosus* has a positive effect on the yield of lactic acid; this was due to the ability of *L. pentosus* to utilize both hexose and pentose sugars. The co-fermentation of *L. casei* and *L. rhamnosus* on rice straw contributed higher lactic acid yield (33.95±0.56g/L) than the co-fermentation with *L. casei* and *L. pentosus.* However, the productivity is slightly lower (0.481 g/L.h) than the CP system. Maximum yield of LA was observed when *L. rhamnosus* was added as thrice the amount of *L. casei* in the fermentation medium. This can be due to the longer duration *L. rhamnosus* needs to reach log phase compared to *L. casei*. Hence, higher amount of *L. rhamnosus* can complement the higher productivity of *L. casei*. It is evident that both the co-fermentation systems (CP and CR) resulted in enhanced LA concentration at the expense of volumetric LA productivity when compared to monoculture systems. Similar trends of simultaneous co-fermentation with LAB were reported previously. For example, Zhang et al., 2015, reported that monocultures of *L. plantarum* and *L. brevis* synthesized LA of concentration 24.3 ± 0.4 g/L (1.01 ± 0.02 g/L.h) and 17.2 ± 0.5 g/L (0.36 ± 0.01 g/L.h), respectively from alkali-treated corn stover with simultaneous enzymatic saccharification and co-fermentation. The co-fermentation enhanced the LA concentration to 28.3 ± 0.2 g/L but with reduced productivity (0.59 ± 0.00 g/L.h) (Zhang and Vadlani, 2015). However, Chen et al., 2020 reported that simultaneous co-fermentation of *L. rhamnosus* and *B. coagulans* resulted in improved LA concentration (112.5 g/L) and LA productivity (2.74 g/L.h) from cassava bagasse compared to monocultures of *L. rhamnosus* (31.0 g/L and 1.94 g/L.h) or *B. coagulans* (30.0 g/L and 1.50 g/L.h) (Chen et al., 2020).

##### 3.4.2.2 Effect of inoculum size

The performance of co-cultures at different inoculum volumes in LA production was studied to find out the sufficient inoculum size required and are graphed in Fig 4e. Within the selected range of 6% −14%, the CP system showed higher LA production at 10% (v/v) and that of CR system was at 8% (v/v). It was found that the increase in inoculum volume from 8% to 10% resulted in an LA concentration change of 27.07±0.62 g/L to 31.74±0.25 g/L. However, inoculum volume beyond 10% (v/v) did not show much changes in the LA concentration. Similar trend was observed in the CR system, where 6%-8% increase in inoculum volume increased the LA production from 27.95±0.62 g/L to 33.42±0.87 g/L. From the study it can be concluded that for the CP system 10% (v/v) inoculum volume and for the CR system 8% inoculum was sufficient to ferment the sugars produced during saccharification. The sigmoid curve for the effect of inoculum size on product formation was observed by several reports where after saturation further changes in inoculum volume was not translated into LA production (Aboseidah et al., 2017; Qi and Yao, 2007).

A set of validation experiments were conducted for CP and CR systems with the best conditions obtained from the preliminary studies. HPLC analysis was conducted to estimate the concentration and purity of LA and is given in Table 3. Similar findings of co-fermentation studies for lactic acid fermentation was reported by Haokok C et al., 2023, where sugarcane bagasse was pretreated by alkali-thermal method followed by saccharification with commercial cellulase and xylanase and co-fermentation by *Lactiplantibacillus plantarum* and *Levilactobacillus brevis* consortia isolated from fermented fish and pork. They reported that the co-fermentation produced 91.9 g/L LA (0.86g/g FS) with a productivity of 0.85 g/L.h whereas the monoculture of *L. plantarum* produced 86.7 ± 0.2 g/L LA with a productivity of 0.8 g/L·h. The comparison of LA production efficiency from co-fermentation and monoculture showed that the former resulted in an improvement of 32% product yield and 14% economic yield (Haokok et al., 2023).

**Table 3.**
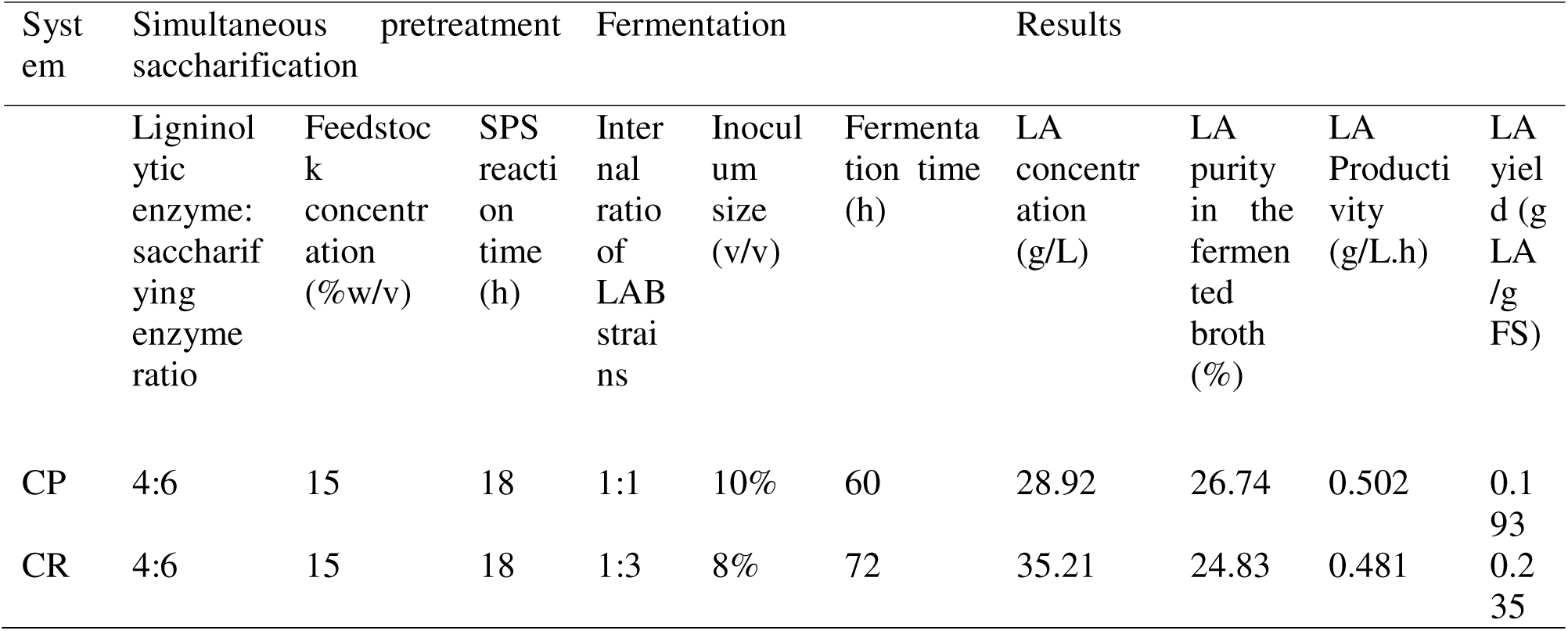
Overview of CP and CR systems and their respective LA concentration, purity, productivity, and yield.

#### 3.4.3 Decolorization and purification of LA

The fermentation broth was decolorized by PAC prior to purification by chromatography. The pigmentation and a fraction of impurities were removed from the broth. The effect of PAC dosage on the color of the fermentation broth and LA recovery was studied, which is shown in Fig 4f. It is evident that 15% PAC could make the broth colorless and suitable for chromatography. The similar trend of decolorization and product recovery was reported by de Oliveira J et al., 2021 where higher PAC loading resulted in low lactic acid yield (de Oliveira et al., 2021). The reduction in LA recovery suggests that in addition to pigment adsorption, PAC was adsorbing LA in their pores, reducing the efficiency of the decolorization process. HPLC analysis of the decolorized sample showed that after 15% PAC treatment the LA concentration in broth from CP system dropped from 28.92g/L to 13.45 g/L with a purity of 80.99% and that of CR system from 35.21g/L to 20.18g/L with a purity of 77.73%. These decolored broths were concentrated to 25g/L LA concentration before subjecting to chromatography. The weak base ion exchange resin Amberlite^®^IRA-67 is a hydrophilic acrylic based resin with high affinity towards lactic acid (Din et al., 2022). The HPLC analysis of purified samples from CP and CR systems reported a 76.77% and 79.66% purity, respectively. The results suggest that for broth from CP system the absorption capacity of the resin was 108.60 mg LA/g resin and that of CR system was 105.24 mg LA/g resin. The absorption capacity of the resin complies with the several reported studies such as 136.11 mg/g as per Zaini, et al 2019 (Zaini et al., 2019) and 150.00 mg/g as per Boonmee, et al 2015 (Boonmee et al., 2016).

A comparison of the two co-fermentation systems (CP and CR systems) shows that the final concentration of lactic acid is a function of growth rates of LAB strains, enzyme activities and substrate utilization rate, and the production of by-products. Hence, an optimization study on inoculation rate, sequential co-fermentation, and time of *L. casei* and *L. rhamnosus* in the co-fermentation was essential to establish a fermentation system to maximize the utilization of hexose and pentose sugars. Thus, from the preliminary experiments, it can be inferred that the CR system is better at yielding lactic acid from rice straw using enzymatic pretreatment and saccharification. Hence, the CR system was further studied using statistical optimization to mathematically model the process.

### 3.5 Central Composite Design for Optimization of LA using CR system

A second-order mathematical model was designed for the CR fermentation system using CCD based RSM. It was noted from preliminary studies that the ability of *L. rhamnosus* to ferment xylose in addition to glucose can be explored by leveraging a sequential inoculation approach. The preliminary studies have shown that there was a decrease in pH of the broth due to LA production. This can disrupt microbial growth and eventually lead to a decline in LA production rate. This kind of product inhibition is handled by regular removal of the product or by introducing neutralizing agents (Rawoof et al., 2021). The study on the CR fermentation system with buffering agent (CaCO_3_) showed better LA concentration in the fermentation broth, sounding the importance of buffering (Supplementary Table S1). Table 4 shows the independent variables, values at their coded levels, and their respective experimental LA concentrations and predicted LA concentrations by the regression equation for the CR system. The quadratic equation generated by regression analysis is expressed in Eq 5. The developed regression model explained the influence of the four independent variables, namely, A-fraction of *L. casei* in the consortium (%), B-Total inoculum volume (v/v%), C-time interval between *L. casei* and *L. rhamnosus* inoculation (h), and D-fermentation time after *L. rhamnosus* inoculation (h).

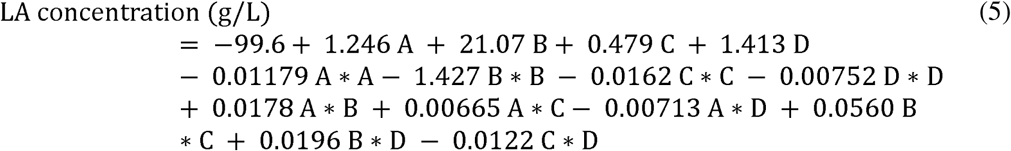

**Table 4.** Design of Experiments (DOE) by CCD for optimization study with the experimental and predicted LA concentration.

| Run no. | A: Fraction of *L. casei* in the consortium (%) | B: Total inoculum volume (v/v%) | C: Time interval between *L. casei* and *L. rhamnosus* inoculation (h) | D: Fermentation time after *L. rhamnosus* inoculation (h) | Experimental LA concentration (g/L) | Predicted LA concentration (g/L) |
| --- | --- | --- | --- | --- | --- | --- |
| 1 | 40 | 6 | 6 | 24 | 40.59±5.97 | 41.81 |
| 2 | 80 | 6 | 6 | 24 | 35.22±1.10 | 34.08 |
| 3 | 40 | 10 | 6 | 24 | 39.49±8.05 | 40.83 |
| 4 | 80 | 10 | 6 | 24 | 33.25±1.95 | 35.94 |
| 5 | 40 | 6 | 18 | 24 | 45.14±3.31 | 46.60 |
| 6 | 80 | 6 | 18 | 24 | 44.27±0.65 | 42.06 |
| 7 | 40 | 10 | 18 | 24 | 50.74±0.58 | 48.30 |
| 8 | 80 | 10 | 18 | 24 | 49.14±1.56 | 46.61 |
| 9 | 40 | 6 | 6 | 48 | 54.14±1.36 | 56.95 |
| 10 | 80 | 6 | 6 | 48 | 40.27±6.95 | 42.37 |
| 11 | 40 | 10 | 6 | 48 | 55.98±2.01 | 57.85 |
| 12 | 80 | 10 | 6 | 48 | 47.30±1.69 | 46.12 |
| 13 | 40 | 6 | 18 | 48 | 61.26±2.60 | 58.22 |
| 14 | 80 | 6 | 18 | 48 | 47.90±0.19 | 46.84 |
| 15 | 40 | 10 | 18 | 48 | 60.39±4.87 | 61.81 |
| 16 | 80 | 10 | 18 | 48 | 54.83±0.78 | 53.27 |
| 17 | 60 | 8 | 12 | 36 | 57.04±4.03 | 59.57 |
| 18 | 60 | 8 | 12 | 36 | 58.83±2.92 | 59.57 |
| 19 | 60 | 8 | 12 | 36 | 62.68±6.69 | 59.57 |
| 20 | 60 | 8 | 12 | 36 | 59.47±5.00 | 59.57 |
| 21 | 20 | 8 | 12 | 36 | 48.45±2.27 | 46.10 |
| 22 | 100 | 8 | 12 | 36 | 27.42±5.78 | 29.83 |
| 23 | 60 | 4 | 12 | 36 | 31.36±1.36 | 31.27 |
| 24 | 60 | 12 | 12 | 36 | 36.55±5.84 | 36.72 |
| 25 | 60 | 8 | 0 | 36 | 53.41±0.71 | 48.53 |
| 26 | 60 | 8 | 24 | 36 | 55.52±5.00 | 60.47 |
| 27 | 60 | 8 | 12 | 12 | 40.82±7.99 | 41.60 |
| 28 | 60 | 8 | 12 | 60 | 64.11±1.95 | 63.40 |
| 29 | 60 | 8 | 12 | 36 | 62.68±4.74 | 56.83 |
| 30 | 60 | 8 | 12 | 36 | 51.57±0.19 | 56.83 |

The second-order polynomial model suggested by regression analysis was examined. The negative constant term (−99.6) in the quadratic equation suggested the validity of the model only within the experimental parameter ranges since it is physically impossible to get a negative value for LA concentration when all the input factors are at zero value. The equation showed that all the linear terms (A, B, C, and D) have a positive linear effect on LA concentration. The negative quadratic effects of the factors (A^2^, B^2^, C^2^, and D^2^) indicated that as the factor increases, the linear positive effect was weakened by the quadratic term which eventually led to response maximum in the selected range. The positive coefficients of the interaction terms A*B, A*C, B*C, and B*D indicated that their proportional increase could optimize the response. Overall analysis of the proposed quadratic model revealed that B: Time interval between *L. casei* and *L. rhamnosus* inoculation was the most influential factor among the four. However, considering the interactions between the factors, an optimization to maximize LA concentration is of utmost relevance. The results of ANOVA analysis of the regression model are described in Table 5. It showed that the model for LA production from CR system is significant due to low p-value (<0.05) for the model and high p-value (>0.05) for lack-of-fit. The significant quadratic coefficients and interaction coefficients are reported in the ANOVA (Table 5); smaller the p-value the more significant the term. It can be estimated from ANOVA (Table 5) that for the quadratic model with DF (degree of freedom) = 17, the f-value (18.29) and probability value (p<0.001) are significant. The high R^2^ value indicates that independent variables are responsible for 88.10% of changes in LA concentration, while only 11.90% was not explained by the regression model. The high value for adjusted R^2^ (83.28%) and predicted R^2^ (75.4%) backs the significance of the model by ANOVA. The closeness of actual experimental LA concentration and the predicted values is visualized in Supplementary Fig S1.

**Table 5.** Analysis of Variance (ANOVA) for CCD-based RSM of one-pot LA production.

| Source | DF<sup>a</sup> | Seq SS | Contribution | Adj SS | Adj MS | F-Value | P-Value |
| --- | --- | --- | --- | --- | --- | --- | --- |
| Model | 17 | 5804.69 | 88.10% | 5804.69 | 341.45 | 18.29 | 0.000 |
| Blocks | 3 | 220.66 | 3.35% | 220.66 | 73.55 | 3.94 | 0.015 |
| Linear | 4 | 2736.49 | 41.53% | 2736.49 | 684.12 | 36.65 | 0.000 |
| Square | 4 | 2671.11 | 40.54% | 2671.11 | 667.78 | 35.77 | 0.000 |
| 2-Way | 6 | 176.43 | 2.68% | 176.43 | 29.40 | 1.58 | 0.178 |
| Interaction |  |  |  |  |  |  |  |
| Error | 42 | 784.05 | 11.90% | 784.05 | 18.67 |  |  |
| Lack-of-Fit | 34 | 601.08 | 9.12% | 601.08 | 17.68 | 0.77 | 0.720 |
| Pure Error | 8 | 182.97 | 2.78% | 182.97 | 22.87 |  |  |
| Total | 59 | 6588.74 | 100.00% |  |  |  |  |

<sup>a</sup>Degrees of freedom; <sup>b</sup>Sum of squares; <sup>c</sup>Mean squares.

Surface plots translate polynomial model obtained from regression analysis to a visual representation of the effect of input variables (independent variables) on the response. The surface plots for each combination of independent variables were generated and shown in Fig 5. The surface plots help to identify the optimum process conditions and sensitivity of the response to the input variables. Fig 5a represents the combined effect of fraction of *L. casei* in the consortium (A) and total inoculum volume (B) on LA production. The interaction surface had taken a parabolic hill shape indicating the selected range of process conditions (A and B) were well-chosen around the optimum. The trend in increase in LA concentration with increase in A and B and decrease beyond the optimum values deduces the rapid sensitivity and robustness of interaction. The upward-opening paraboloid shapes of Fig 5b, c, d, and e indicate that the optimum conditions for maximum LA concentration lie well within the experimental range of input parameters and the response increases towards the center of the design region. The ridge region of 3D plots represents the maximum LA concentration in the experimental range and it shows the region of least change where the input parameters are less sensitive towards the response. The interaction between A and C (Fig 5b) showed that the ridge region was between 40-60% of *L. casei* in the consortium and >15h inoculation gap between *L. casei* and *L. rhamnosus*. This can be deduced that anything beyond these values of A and C can either result in underutilizing *L. casei* for hexose fermentation or overutilizing *L. rhamnosus* for hexose and pentose fermentation and thereby risking the productivity of the process. The wide titled ridge region of A and D interaction (Fig 5c) indicates a positive interaction between the two parameters and there is a possibility of substantial tradeoffs between them. The ridge is prominent between 20% <A<70% and D>40 h which indicates without *L. rhamnosus* in the system for a defined amount of time the yield and productivity of the process drop making the system infeasible. The narrow ridge of B and C interaction plots (Fig 5d) notify the low robustness and high sensitivity in the ridge direction. The narrow range of B (7%<B<9%) and C (>15 h) infers that increasing the total inoculum volume beyond the range do not promise high LA yield and in fact, it might decrease the LA concentration due the increased demand of carbon source for microbial growth at high inoculum volumes restricting the LA production metabolism. The ridge of B (7%>B>10%) and D (>40h) interaction (Fig 5e) shows a robustness in experimental range for better LA production. However, a higher or lower inoculum volume can either result in a low-density fermentation system or a substrate-limited system adversely affecting the efficiency of the process. The ridge in the interaction of C and D (Fig 5f) shows a high-sensitivity optimum in the experimental range. It indicates that the system prefers longer time intervals between *L. casei* and *L. rhamnosus* inoculation (15h<C<20h) and longer fermentation after *L. rhamnosus* inoculation (D>50h) for better credentials. The inferences from contour plots of interactions (Supplementary Fig S2) align with the interpretations of 3D interaction plots.

**Fig 5.**
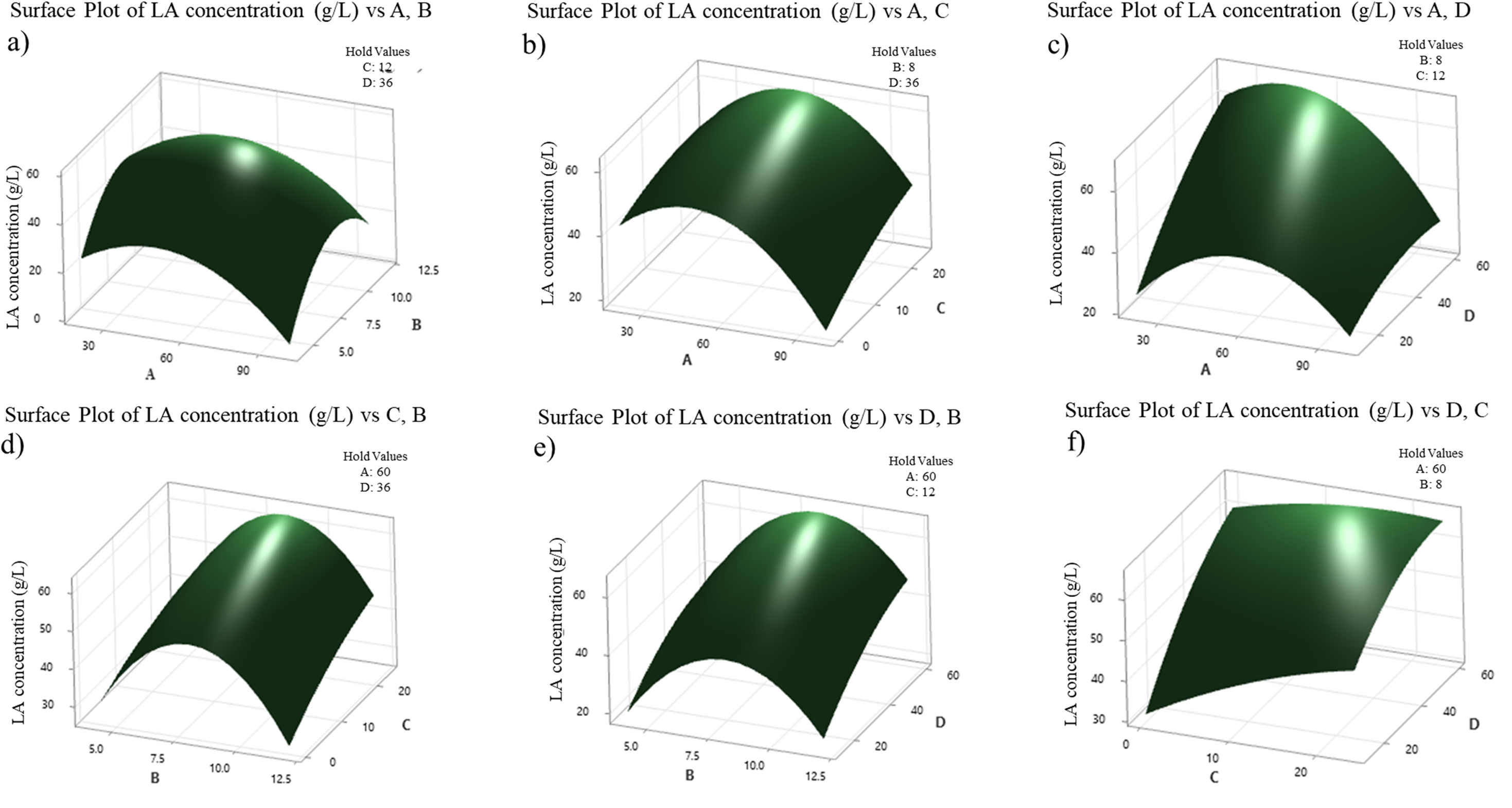
3D plots characterizing the interactions of fermentation conditions on LA concentration: a) A and B, b) A and C, c) A and D, d) C and B, e) D and B, and f) D and C

#### 3.5.1 Optimization and Validation of the model

The predicted regression model was optimized to maximize the LA concentration from the experiment, and the optimum values for four input variables were estimated. The predicted model was validated by conducting experiments at the predicted optimum input variables; fraction of *L. casei* in the consortium (A)-45.8%, total inoculum volume (B) – 8.4% (v/v), Time interval between *L. casei* and *L. rhamnosus* inoculation-16h, and Fermentation time after *L. rhamnosus* inoculation-60h. The validation experiment showed 64.96 g/L which was closer to the predicted LA concentration, 68.12 g/L. The yield and productivity of LA from the validation experiment was estimated to be 0.316 g LA/g rice straw (corresponding to 0.74±0.04 g/g initial reducing sugars) and 0.855 g/L.h, respectively. Tu et al., 2019 reported comparable lactic acid production (65.6 g/L and 0.38g LA/g rice straw) in a simultaneous saccharification and fermentation setup using commercial cellulase enzyme followed by *L. plantarum* fermentation on hydrolysate from acid pretreatment of rice straw (Tu WeiLin et al., 2019). In a similar study by Chen et al., 2019, LA was produced from rice straw by dilute ethylenediamine pretreatment followed by fed-batch simultaneous saccharification and fermentation using *B. coagulans*. The process produced 92.5 g/L LA with a yield of 0.272g LA/g rice straw (Chen et al., 2019). Similar results were reported by Sivagurunathan et al., 2022 for an LA production system employing mild acid pretreatment, enzymatic saccharification and fermentation with *L. lactis* and rice straw as feedstock (82.2 □±□1.9 g/L LA and 0.411g LA/g rice straw) (Sivagurunathan et al., 2022). The improved LA yield with respect to rice straw consumption in the present study can be elucidated as a result of the one-pot fermentation process where the reduction of unit operations has reduced the loss of raw material for fermentation. Ouyang et al., 2020 reported LA production using wheat straw which was subjected to dilute sulphuric acid pretreatment followed by simultaneous enzymatic saccharification and fermentation of *Bacillus coagulans*. The process produced 26.30 g/L LA (0.252 g/L.h) with 70.9% yield on holocellulose basis (0.444g/g feedstock) (Ouyang et al., 2020). On the other hand, Shahab et al., 2018 reported a consolidated bioprocessing system for LA production from steam pretreated washed Beech wood using a fungal-bacterial consortium for simultaneous saccharification and fermentation. They employed *Trichoderma reesei* and *L. pentosus* for saccharification and fermentation, respectively and produced 19.8 g/L LA with a productivity of 0.1g/L.h in a solid-state fermentation condition (Shahab et al., 2018). These studies enlighten the possibility of reducing unit operations and omitting the need for inter-process material transfer without compromising the product yield. In the present study, the employment of sequential fermentation where the LA fermentation efficiency of *L. casei* was utilized maximum before introducing *L. rhamnosus* had shown to improve the efficiency of the system. It was observed that initially (at 0 h of fermentation), the fermentation media contained hexose sugars and pentose sugars in concentrations, 56.83g/L and 43.96g/L respectively. After 16h of fermentation (before *L. rhamnosus* inoculation), the concentration declined to 33.15g/L and 41.83 g/L and later declined to 0.05g/L and 25.54g/L at the end of fermentation (60h after *L. rhamnosus* inoculation). This interprets that approximately 25.59% of total sugars (hexose and pentose) were utilized by *L. casei* in the first 16h of fermentation and the *L. casei-L. rhamnosus* consortium was able to consume 65.88% of total sugar content. Whereas the CR system with sequential fermentation consumed 74.61% of total initial sugar content corresponding to 75.19 g/L of RS. In which hexose sugar consumption was 99.9% and pentose sugar consumption was 41.89% making the yield of LA 0.6445g/g initial fermentable sugars. This validates the hypothesis that *L. rhamnosus* is able to metabolize pentose sugars (Eg. Xylose) only in the absence of hexose (glucose) sugars. Hence, it can be inferred that sequential co-fermentation results in a better fermentation system if designed in the order of metabolic preference of the microorganisms involved. Qiu et al., 2022 reported the production of D-LA using *Pediococcus acidilactici* from undetoxified acid-pretreated corncob slurry. They reported that since the undetoxified slurry contained xylose sugars and aldehyde inhibitors, to bypass the competitive inhibition of cellulase enzyme, *P. acidilactici* was inoculated first followed by cellulase (after 24h) for saccharification. *P. acidilactici* is known to ferment both xylose and glucose. This has improved the LA production from 39.2 g/L (simultaneous addition of cellulase and *P. acidilactici*) to 61.9 g/L LA (sequential addition of *P. acidilactici* and cellulase) since *P. acidilactici* is known to ferment both xylose and glucose sugars (Qiu et al., 2022). The study elucidated the importance of order of unit operations specific for each bioprocess and the advantages of consolidated bioprocessing systems for better utilization of feedstock components.

The higher productivity of the co-fermentation system in the optimized process in comparison with the unoptimized system (in section 3.4.2) can be interpreted as the result of addition of CaCO_3_ as buffering agent. It was reported that certain calcium salts convert lactic acid into lactate salts thereby preventing product inhibition in the system by maintaining the pH of the broth allowing the microorganisms to sustain in the fermentation media longer (Zhang and Vadlani, 2013). In addition, a time interval in the inoculation between the two strains aided LA production in such a way that it maximized the utilization of hexose sugars followed by pentose sugars by providing adequate time for *L. casei* to synthesize LA and leave enough glucose for *L. rhamnosus* to thrive through its lag and exponential phases of growth. Thus, it can be inferred that 45.8% of *L. casei* and 16h co-fermentation interval balanced cell growth and product synthesis by optimizing nutrient competition and product inhibition. It can be noted that the final LA concentration in the fermentation broth is a function of initial fermentable sugars and the buffering capacity of the broth. The results from a comparison study on simultaneous and sequential inoculation of two strains of LAB; *E. casseliflavus* and *L. casei* for LA production resembles the results of the present study. Where, an inoculation gap of 49h resulted in 95g/L LA (0.63g/g) whereas the simultaneous inoculation produced 70g/L (0.46g/g) (Taniguchi et al., 2004). However, on the contrary, a sequential inoculation did not benefit Cui et al., 2011 in terms of LA production or yield when *L. rhamnosus* and *L. brevis* were used. They reported that simultaneous inoculation resulted in 14.80 g/L LA (0.73 g/g) and sequential inoculation when *L. brevis* was added after 12h of *L. rhamnosus* inoculation resulted in 13.44 g/L LA (0.72g/g). Whereas the monoculture fermentations produced similar range of LA for *L. rhamnosus* (13.73 g/L and 0.79g/g) and *L. brevis* (13.44g/L and 0.67g/g). Hence, the inoculation strategy and selection of LAB strains play an essential role in improving the credentials of a fermentation system. Only fewer studies are reported on the one-pot fermentation approach for LA production. Yadav N et al., 2021 reported an LA yield of 36.75 g/L (0.612 g/g substrate) with 0.51 g/L.h productivity using *L. plantarum* SKL-22 strain for fermentation of ionic liquid (1-Ethyl-3-methylimidazolium-acetate) pretreated-cellulase saccharified rice straw (Yadav et al., 2021). Similarly, Grewal and Khare 2018 reported a one-pot system with ionic liquid (1-Ethyl-3-methylimidazolium-acetate) pretreatment, nanoimmobilized cellulase for saccharification, and *L. brevis* for fermentation using sugarcane bagasse as raw material. The system produced 0.52 g LA/g from sugarcane bagasse with simultaneous consumption of hexose and pentose sugars during fermentation by *L. brevis* (Grewal and Khare, 2018). However, there is a need for stability improvement of enzymes and microbial strains prior to fermentation when chemical pretreatments are employed. This can complex the upstream processing of the system due to the time-intensive processes and the possibility of genetic instability in future. Thus, it can be interpreted that the present work proposes a feasible one pot fermentation strategy for LA production from rice straw with a complete biological route. The process omitted inter-process material transfer between pretreatment, saccharification, and fermentation reducing the chances of material loss and energy consumption.

### 3.6 Downstream processing of LA

HPLC analysis of samples after fermentation, decolorization, and purification is shown in Fig 6a. The analysis revealed that the one-pot co-fermentation system produced 64.96 g/L LA with a purity of 28.56% and after PAC decolorization the purity improved to 68.21%. Meanwhile, the ion exchange chromatographic purification has enabled LA broth with 85.56% purity which complies with the industrial standard of LA in plastic industries (50-80% purity) (Vijayakumar et al., 2008). The overall efficiency of the DSP was estimated to be 45.29%. In which PAC decolorization recovered 66.67% LA and the chromatographic purification by Amberlite^®^IRA-67 resin showed 68.36% recovery. Similar results were reported by Bernardo et al., 2016, where the LA fermentation broth produced from dairy industry waste using *L. rhamnosus* B103 in fed batch operation yielded 143.7 g/L LA. The DSP of the system recovered 10.30% LA with 63.70% purity. Here, the PAC decolorization (18.72% w/v) showed 77.00% recovery and that of purification by Amberlite^®^IRA-67 resin chromatography was 13.4% (Bernardo et al., 2016). Whereas Zaini et al., 2019 reported an overall LA recovery of 80.4% after PAC decolorization (7% w/v) and purification by chromatography using Amberlite^®^IRA-67 resin. The system was designed to synthesize LA from dried distillers’ grains (DDGS) as feedstock using *L*. *coryniformis* subsp. *torquens* via alkali pretreatment and simultaneous saccharification and fermentation which produced 25.9g/L LA (Zaini et al., 2019). Further, Ahmad et al., 2021 reported a downstream system with PAC decolorization (5% w/v) and chromatography by Amberlite^®^IRA-67 was able to recover 91% LA with 94.6% optical purity. The fermentation system used date pulp waste as feedstock and indigenous microbiota for fermentation (Ahmad et al., 2021). These studies showed that the decolorization step largely influences the overall recovery and efficiency of DSP of LA production systems. The color of the fermented broth is an outcome of raw materials added at pretreatment, saccharification, and fermentation stages and it is crucial to decolor the broth prior to injecting to chromatography column.

**Fig 6.**
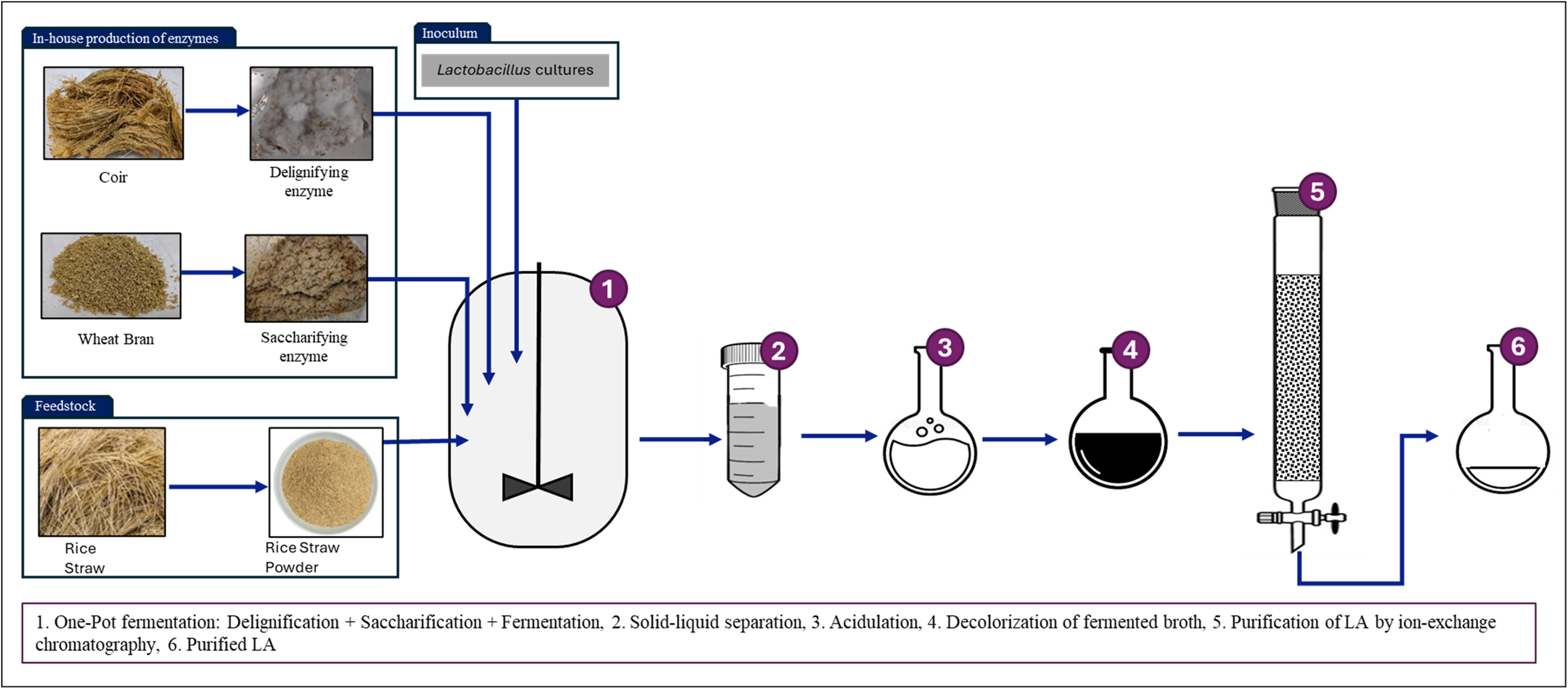
Characterization of the recovered LA: a) Purity of LA samples at different stages of fermentation-purification and b) optical purity of LA extracted from the process.

CD spectroscopy was used to assess the optical purity of lactic acid after purification by chromatography. The CD spectra of the sample was compared with that of standard (L-LA) as shown in Fig 6b and the optical purity was determined by comparing the molar ellipticity at 210nm (Eq 4). It was estimated that 84.95% optical purity was obtained for the produced lactic acid. The purity indicates that the purified broth contains 84.95% of L-LA which can be advantageous for applications such as synthesis of polylactic acid (PLA) with amorphous structure, enhanced flexibility, and faster biodegradation. The optical purity of the fermented broth is a function of the microbial strains involved in the consortia, type of fermentable sugars released during saccharification, microbial mechanisms activated during fermentation, and the purification strategies in DSP. López-Gómez et al., 2020 reported that incorporating a sequential purification by microfiltration, nanofiltration, and electrodialysis improved the optical purity of LA in fermentation broth from 93% to 98.7% when LA was produced from hydrolysate of municipal solid waste (López-Gómez et al., 2020a). Similarly, Olszewska-Widdrat et al., 2019 reported 98.9% optically pure LA after purification of *B. coagulans* fermented broth of sweet sorghum juice when coarse and ultra-filtration, electrodialysis, chromatography by Amberlite^®^IRA-92 and distillation was sequentially employed in the DSP (Olszewska-Widdrat et al., 2019). Hence, further purification strategies can be incorporated into the DSP to improve the optical purity of LA. It is important to note that optical purity plays a significant role in determining the fate of LA thus it is essential to purify the fermentation broth considering the intended application and its degree of optical purity requirement.

### 3.7 Material balance and process comparison

Material balance of the one-pot co-fermentation process along with complete downstream processing of LA production is depicted in Fig 7. The one-pot fermentation would be able to produce 316.14g LA from 1000g rice straw. The DSP of the broth would be able to recover 144.08g LA, making the overall LA yield as 0.144g/g rice straw. Table 6 compares the current state of LA research with the present study. Most of the reported studies were based on separate hydrolysis and fermentation (SHF) or simultaneous saccharification and fermentation (SmSF) with already pretreated raw material. In case of SHF, high operating costs and process complexity limit its feasibility. Whereas for SmSF, even though the synergistic activities of saccharifying enzymes and LAB strains were advantageous, the need for LAB strains to tolerate optimal temperature (∼50), and pH (4.8-5.5), and the need for starter sugar content for their initial growth often hurdles the LA yield (Ojo and de Smidt, 2023; Zhang et al., 2022). The credentials of LA for the novel one-pot co-fermentation were comparable to the recently reported studies, and it showed that the strategy developed is relevant in the present scenario. The use of second-generation feedstocks (wheatbran, coir, and rice straw) at both the stages of LA production, namely, SPS and co-fermentation adds the overall carbon credit of the process. The choice of pretreatment and its efficiency is crucial in a one-pot strategy. Over chemical technology, processes involving enzyme technology tend to produce less by-products improving the product quality, impart high specificity enhancing the overall yield and efficiency of the process and milder reaction conditions make the process align towards sustainable development goals (SGDs). Enzyme technology combined with one-pot fermentation can reduce the capital cost, steps, and waste generation in a system. Since all the treatments were conducted in the same vessel, the water usage and sugar loss are reduced aiding productivity, and time and energy efficiency. Along with the green perspectives of fermentation, the DSP developed for the process also backs the sustainability vision. As discussed in section 3.5.1, the present study outsmarts previously reported one-pot studies on LA production (Grewal and Khare, 2018; Yadav et al., 2021)with green motives in terms of yield and productivity.

**Fig 7.**
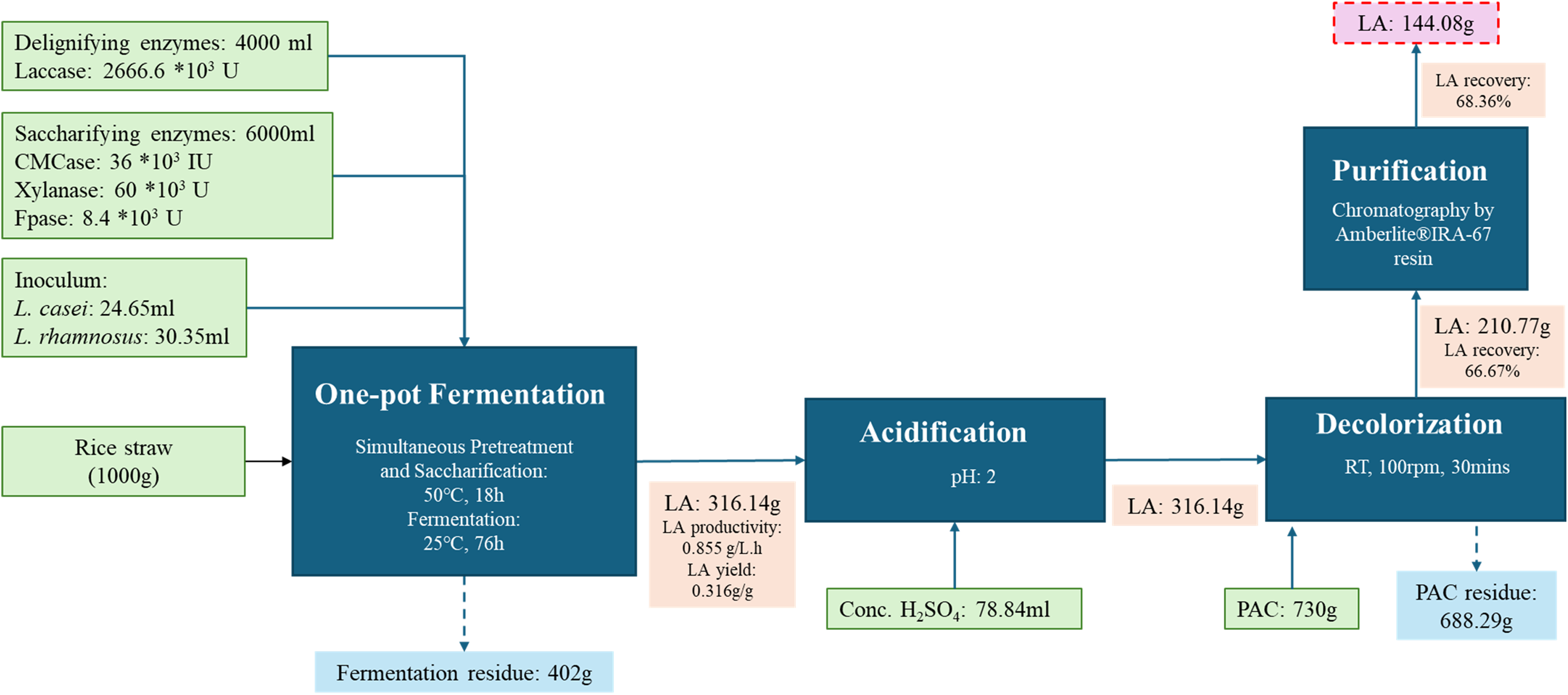
Material balance of the novel one-pot co-fermentation followed by downstream processing of LA production channel from rice straw

**Table 6.**
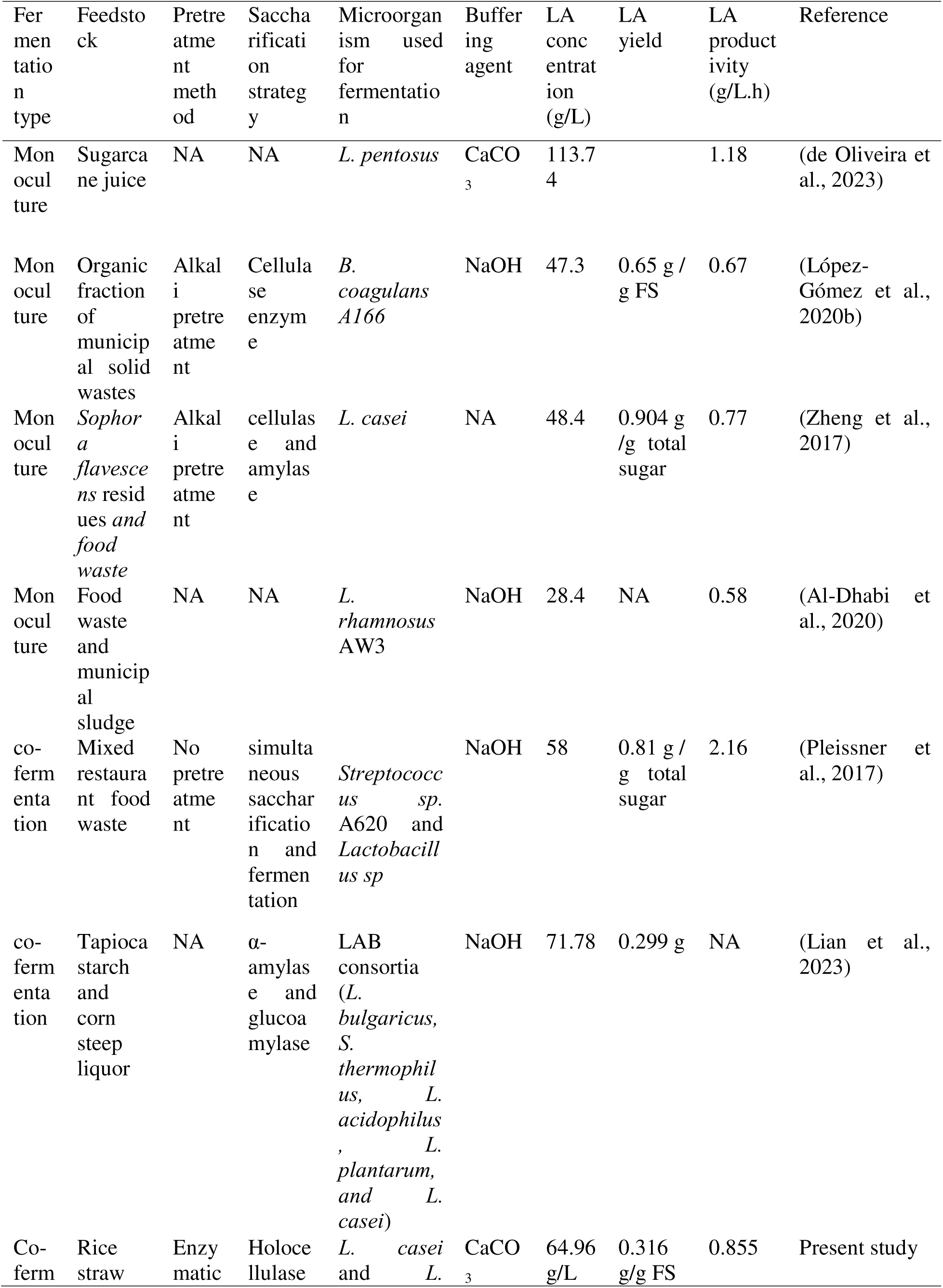
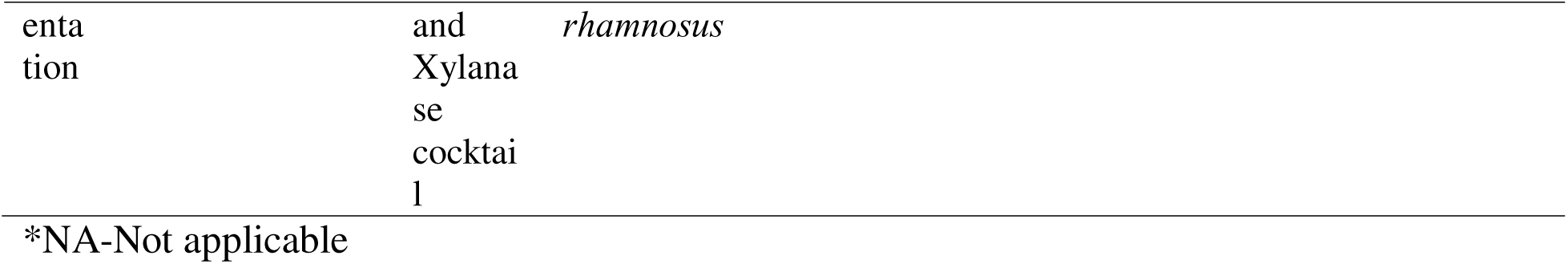
Current advances of lactic acid production from lignocellulosic biomass.

## 4. Conclusions

The present study illustrated the influence of co-fermentation on the yield of lactic acid (LA) via one-pot fermentation approach. An analysis on the production of lactic acid from rice straw reveals that the final lactic acid concentration in the fermentation broth depends on the amount of fermentable sugar produced during enzymatic saccharification, microbial strains involved, and the buffering capacity of the media during fermentation. The use of microbial consortia was found to be effective in utilizing maximal fermentable sugars. After preliminary evaluations of the compatibility of three different lactobacillus strains (*L. casei, L. pentosus* and *L. rhamnosus*), a statistical optimization was conducted with *L. casei* and *L. rhamnosus* co-fermentation system employing one-pot lactic acid production strategy. The optimized system was able to produce 64.96 g/L LA with a productivity of 0.855 g/L.h and yield of 0.6445 g LA/g initial fermentable sugars. Decolorization followed by chromatographic purification of the fermented broth using Amberlite^®^IRA-67 resin yielded a colorless broth with 85.56% purity containing 84.95% optically pure L-LA. The incorporation of fungal enzymes for both pretreatment and saccharification and sequential fermentation strategy to minimize fermentable sugar loss make the process innovative. This novel one-pot lactic acid fermentation approach enhanced the lactic acid production from rice straw in comparison to the conventional fermentation. In addition to the use of in-house enzymes from *P. ostreatus* and *T. reesei,* the downstream processing with PAC decolorization followed by purification by chromatography makes the process move towards sustainability and green manufacturing.

## Author’s contribution

Ms. Bhavya Surendran V S: writing-original draft, data curation, methodology, investigation. Dr. Althuri Avanthi: visualization, supervision, project administration, resources, writing-review and editing. All authors reviewed the manuscript.

## Acknowledgement

This work was supported by Ministry of Education (MoE), India fellowship and ANRF India (SERB-EEQ/BT/F304/2023-24/G667). Authors acknowledge the Director, Indian Institutes of Technology (IIT), Hyderabad for providing other necessary facilities.

## Data availability

Data will be made available on request.

---

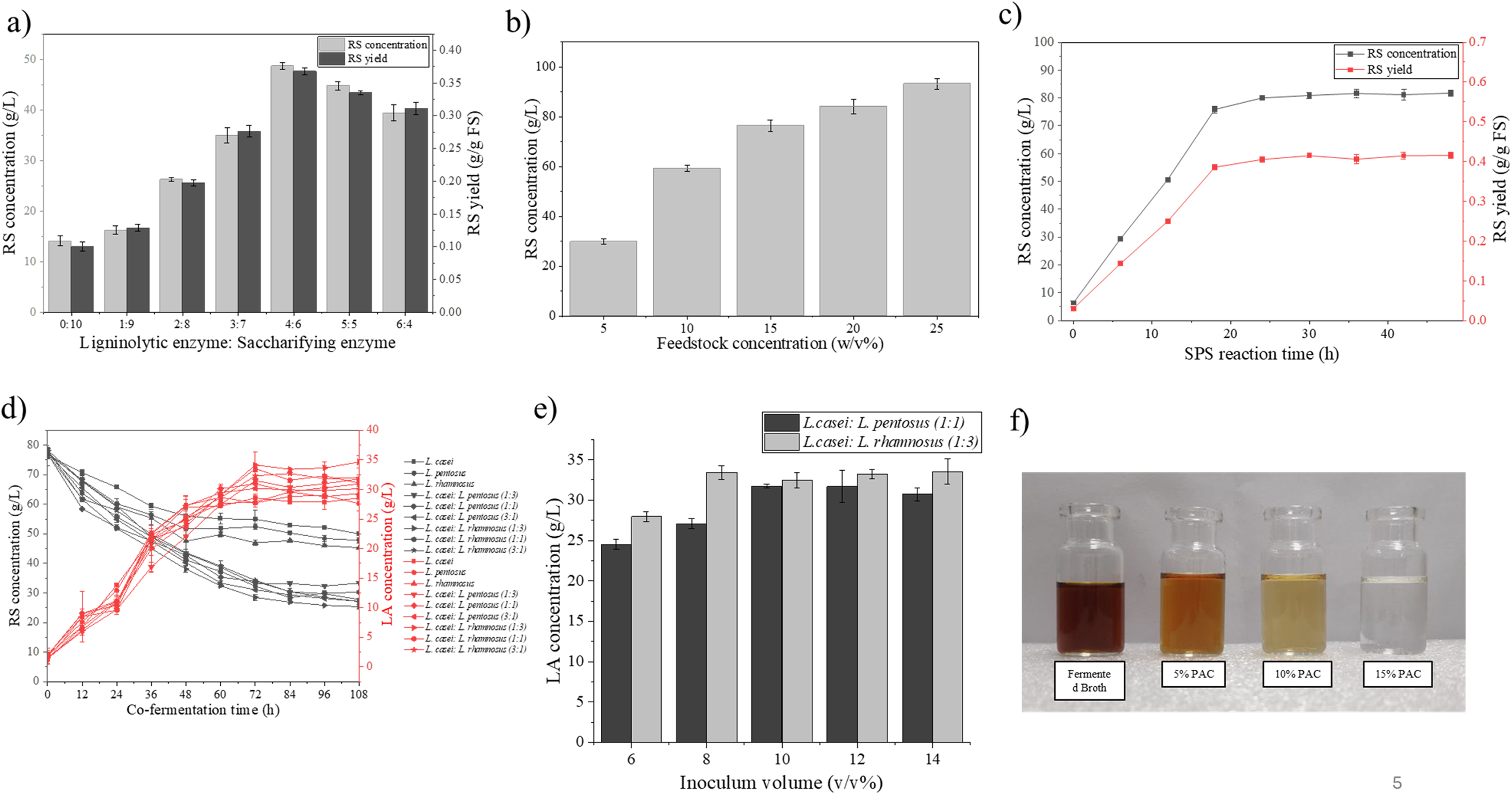

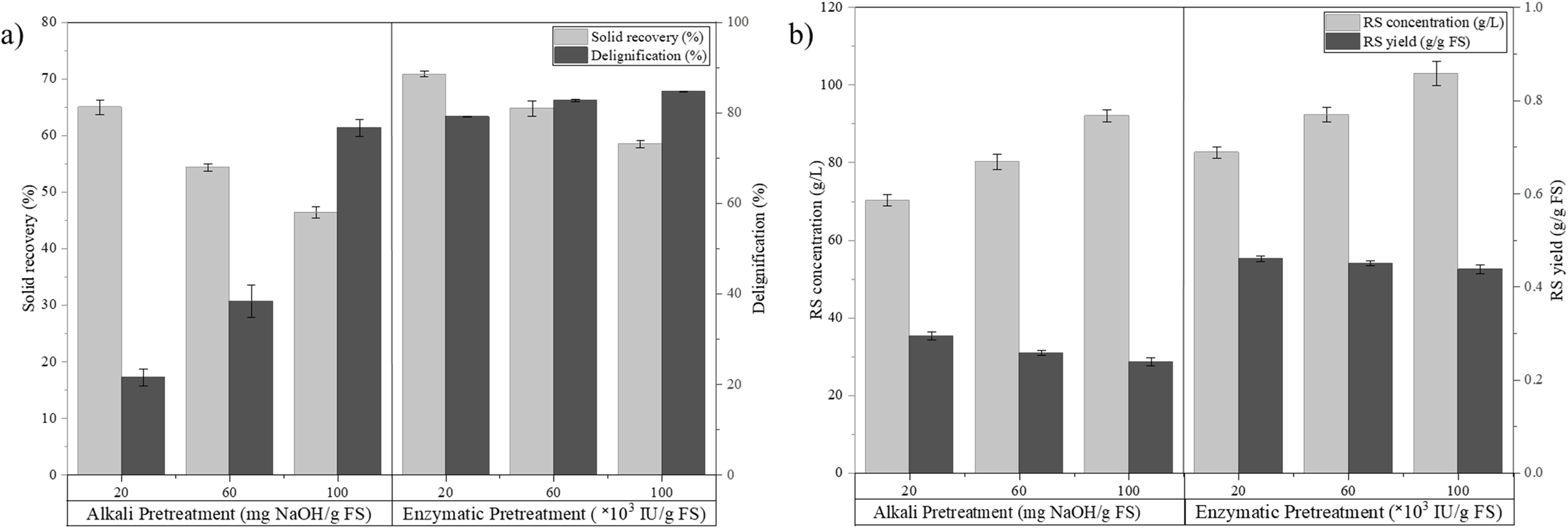

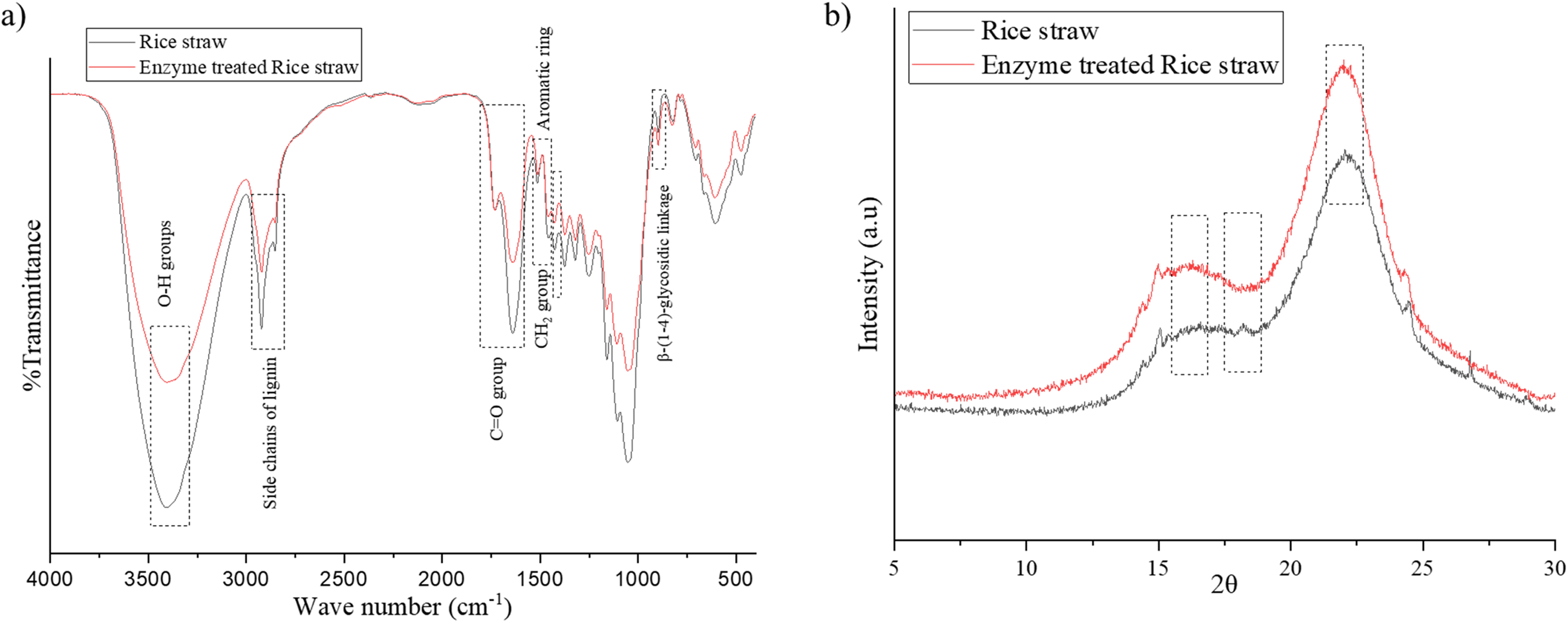

