## Supplementary material for "One-pot lactic acid production from rice straw: A consolidated bioprocess with enzymatic pretreatment-saccharification and Microbial co-fermentation"

Table S1. Effect of CaCO3 concentration on the pH and LA production of CR system.

| CaCO3 Conc (w/v%) | Initial pH | Final pH | LA concentration (g/L) |
| --- | --- | --- | --- |
| 0 | 6.22±0.04 | 3.75±0.01 | 24.42±0.12 |
| 0.1 | 6.38±0.03 | 3.82±0.01 | 40.29±1.50 |
| 0.25 | 6.41±0.02 | 4.375±0.01 | 45.50±0.50 |
| 0.5 | 6.41±0.01 | 5.04±0.01 | 50.12±0.69 |
| 0.75 | 6.47±0.01 | 5.125±0.01 | 50.87±0.37 |
| 1 | 6.53±0.02 | 5.16±0.01 | 52.29±1.37 |

Fig S1. Correlation between actual and predicted values of LA concentration by the regression model

Fig S2. Contour plots characterizing the interactions of fermentation conditions on LA concentration: a) A and B, b) A and C, c) A and D, d) C and B, e) D and B, and f) D and C
